# Gbx2 identifies two amacrine cell subtypes with distinct molecular, morphological, and physiological properties

**DOI:** 10.1101/2020.05.19.104307

**Authors:** Patrick C. Kerstein, Joseph Leffler, Benjamin Sivyer, W. Rowland Taylor, Kevin M. Wright

## Abstract

Our understanding of how the nervous sytem works is limited by our ability to identify the neuronal subtypes that comprise functional circuits. Using a genetic approach, we show that the transcription factor *Gbx2* labels two amacrine cell (AC) subtypes in the mouse retina that have distinct morphological, physiological, and molecular properties. One subtype of Gbx2+ ACs are likely the previously characterized On-type GABAergic CRH-1 AC. The other Gbx2+ AC population is a previously uncharacterized non-GABAergic, non-Glycinergic (nGnG) AC subtype. Gbx2+ nGnG ACs are On-Off type cells with asymmetric dendritic arbors. Gbx2+ nGnG ACs also exhibit tracer coupling to bipolar cells (BCs) through gap junctions that are modulated by dopamine signaling. This study genetically identifies a previously uncharacterized AC subtype and reveals an unusual AC-BC connectivity through gap junctions that may provide a novel model of synaptic communication and visual circuit function.

## INTRODUCTION

The mammalian nervous system is comprised of hundreds of distinct neuronal subtypes that form precise connections with one another. Neuronal subtypes can be defined by a combination of their morphological, physiological, and molecular properties (Zeng & Sanes, 2017). Recent single cell RNA sequencing (scRNAseq) approaches have greatly expanded the catalogue of neuronal subtypes based on transcriptional profiles (Macosko *et al.*, 2015; Saunders *et al.*, 2018; Tasic *et al.*, 2018). However, linking the morphological and physiological properties of neuronal subtypes to their molecular profile and identifying their function within neural circuits remains a major challenge.

The retina is an ideal system to address such questions. It contains a complete neural circuit organized in a highly stereotyped manner within a compact space. Three classes of excitatory neurons—photoreceptors, bipolar cells (BCs), and retinal ganglion cells (RGCs) — connect in sequence to sense light and transmit this sensory information to the brain. Two classes of inhibitory neurons—horizontal cells and amacrine cells — increase the feature selectivity of these sensory signals by providing spatial and temporal regulation of excitatory cell activity (Diamond, 2017). Within these 5 classes, there are over 120 distinct neuronal subtypes (Macosko *et al.*, 2015; Sanes & Masland, 2015; Shekhar *et al.*, 2016; Rheaume *et al.*, 2018; Tran *et al.*, 2019; Yan *et al.*, 2020). This high level of diversity reflects the enormous amount of computation necessary to encode up to 40 distinct representations of the visual field (Baden *et al.*, 2016).

Amacrine cells (ACs) exhibit the greatest diversity in number and variance between subtypes. Morphological analysis of ACs predicts there are ∼45 AC subtypes (MacNeil *et al.*, 1999; Badea & Nathans, 2004; Lin & Masland, 2006; Helmstaedter *et al.*, 2013) while recent single cell transcriptomic analysis predicts more than 60 distinct AC types (Peng *et al.*, 2019; Yan *et al.*, 2020). The data from these scRNAseq studies can provide potential markers for identifying neuronal subtypes in mouse (Macosko *et al.*, 2015; Shekhar *et al.*, 2016; Rheaume *et al.*, 2018; Tran *et al.*, 2019) and primate retinas (Peng *et al.*, 2019).

AC subtypes display characteristic specializations evident in their selective synaptic connectivity and neurotransmitter release. The dendritic morphology and stratification of an AC subtype determines its receptive field size and dictates the potential pre- and post-synaptic partners within the inner plexiform layer (MacNeil & Masland, 1998; Diamond, 2017). Two broad groups of AC subtypes are defined by their expression of either glycine or GABA. In addition to the inhibitory neurotransmitter, some AC subtypes co-release an excitatory neurotransmitter, for example, glycine and glutamate (Haverkamp & Wassle, 2004; Johnson *et al.*, 2004; Lee *et al.*, 2014) or GABA and acetycholine (Brecha *et al.*, 1988; Vaney & Young, 1988). Other ACs also release neuromodulators like dopamine (Newkirk *et al.*, 2013) or neuropeptides (Zalutsky & Miller, 1990). Furthermore, in addition to neurochemical signaling, AC subtypes can form electrical synapses via gap junctions with BCs, RGCs, and other ACs (Vaney & Weiler, 2000; Bloomfield & Volgyi, 2009). Despite this broad functional and morphological diversity, most AC subtypes have not been thoroughly characterized due to a lack of genetic tools to prospectively identify and manipulate them.

Here we identify two AC subtypes that are genetically labeled by a mouse line expressing tamoxifen-inducible *Cre* recombinase from the endogenous locus of the transcription factor *Gbx2* (*Gbx2*^*CreERT2-IRES-EGFP*^). These two AC subtypes have distinct morphological, physiological, and molecular properties. S5-Gbx2+ ACs have larger dendritic arbors that stratify in S5, are exclusively On-type, and are GABAergic. In contrast, S3-Gbx2+ ACs have smaller, dense, asymmetric dendritic arbors that stratify in S3, are On-Off type, and are non-GABAergic non-Glycinergic (nGnG) ACs. S3-Gbx2+ AC subtypes also exhibit an unusual pattern of exclusively heterotypic tracer coupling to BCs, suggesting they communicate in part through electrical synapses. The identification and analysis of the Gbx2+ AC subtypes increases our understanding retinal circuits and vision.

## RESULTS

### *Gbx2*^*CreERT2-IRES-EGFP*^ marks two distinct amacrine cell subtypes

To begin unraveling the neuronal subtype complexity in the retina, we sought to identify *Cre* or *CreERT2* mouse lines that could be used to selectively label and manipulate single neuronal subtypes in the retina. Using scRNAseq datasets and transgenic mouse databases as a guide (Siegert *et al.*, 2009; Macosko *et al.*, 2015), we identified *Gbx2*^*CreERT2-IRES-EGFP*^ as a mouse line predicted to label a sparse population of neurons in the retina (Chen *et al.*, 2009). Since the *CreERT2-IRES-EGFP* cassette is knocked into the *Gbx2* locus, labeled neurons are expected to faithfully recapitulate its endogenous expression pattern. Crossing the *Gbx2*^*CreERT2-IRES-EGFP*^ line to multiple Cre-dependent reporter lines labeled neurons in the inner nuclear layer (INL, Fig. 1a) and the ganglion cell layer (GCL, Fig. 1b). Low doses of tamoxifen (<0.02mg) labeled neurons in both the INL and GCL with dendrites that selectively stratify in sublamina 3 (S3) of the inner plexiform layer (IPL). When we used a saturating dose of tamoxifen (2.0 mg/day for 2 days, Fig. S1d-i), we observed a second neuronal subtype with dendrites that stratify in sublamina 5 (S5) of the IPL (Fig. 1c, g). Importantly, the same two neuronal populations were labeled whether tamoxifen was administered at embryonic (E16), postnatal (P0-2), or adult (P28) ages (Fig. 1 – supplement 1a-c), demonstrating that the *Gbx2*^*CreERT2-IRES-EGFP*^ line provides a consistent and stable method for labeling the same neuronal populations from embryonic to adult stages.

**Figure 1.**
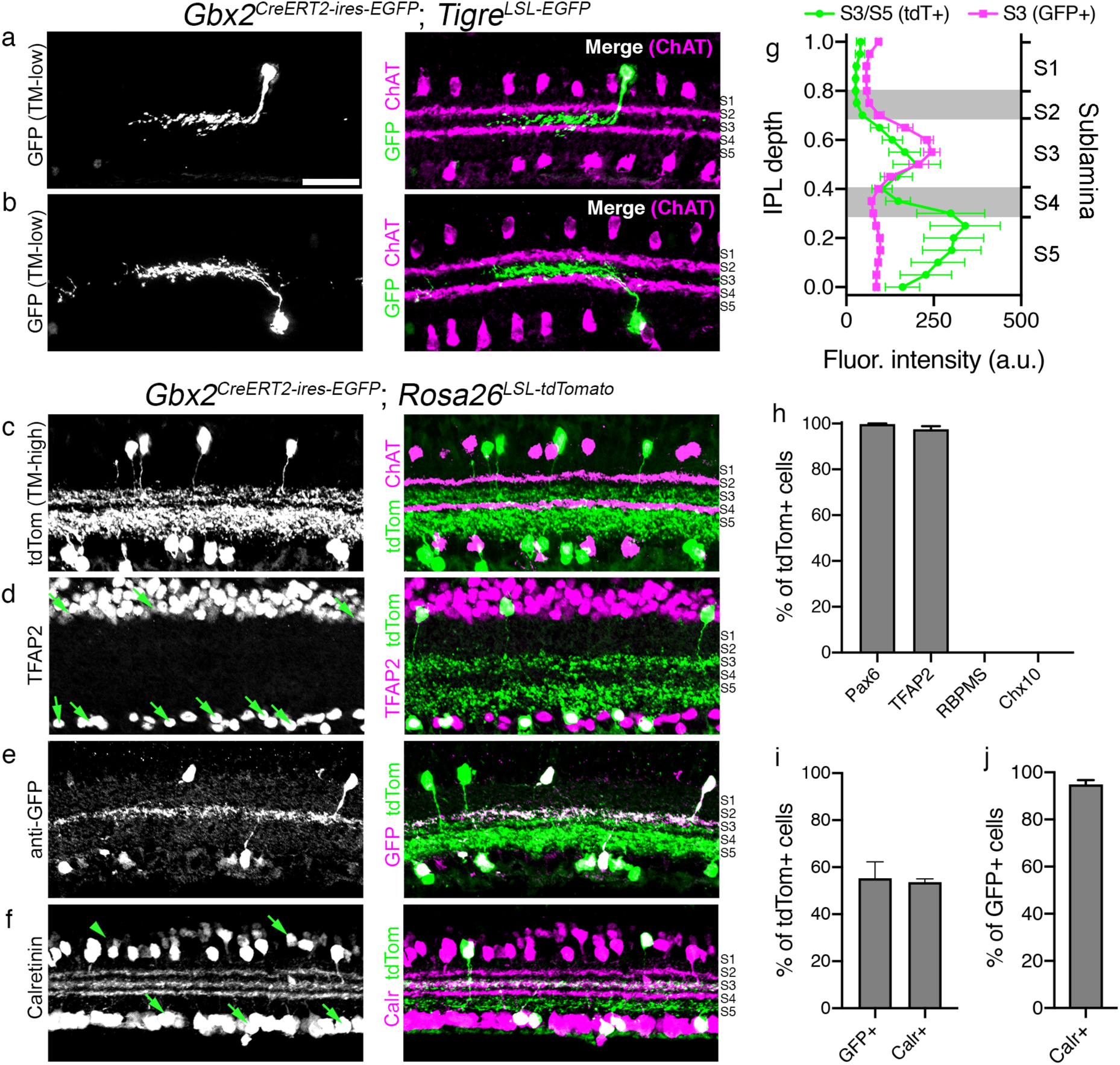
*Gbx2*^*CreERT2-IRES-EGFP*^ selectively labels a subtype of amacrine cells in the mouse retina. (**a-b**) Left: Cross sections of adult retinas from *Gbx2*^*CreERT2-IRES-EGFP*^; *TIGRE*^*TIT2L-GFP-ICL-tTA2*^ mouse labeling single Gbx2+ ACs (low-TM, 0.02mg tamoxifen) in the (**a**) INL and (**b**) GCL. Right: Merged image, ChAT (magenta, S2, S4), Gbx2-GFP (green, S3). (**c**) Cross-sections of an adult retina from a *Gbx2*^*CreERT2*^; *Rosa26*^*LSL-tdTomato*^ mouse labeling the total Gbx2+ AC population (high-TM, 2.0mg tamoxifen). Right: Merged image, ChAT (magenta) and Gbx2-tdTom (green, S3 and S5). (**d-f**) Left, retinal sections from a *Gbx2*^*CreERT2-IRES-EGFP*^; *Rosa26*^*LSL-tdTomato*^ mouse immunolabeled with (**d**) an amacrine cell marker TFAP2, (**e**) GFP, (**f**) calretinin co-labeled with Gbx2+ tdTomato+ retinal neurons (green, right). (**g**) Fluorescent intensity plotted in IPL depth using ChAT bands in S2 and S4 as reference points. Magenta: Gbx2+ ACs detected by EGFP immunohistochemistry (n=6 measurements, 3 retinas); Green: Gbx2+ ACs detected by tdTomato labeling (n=6 measurements, 3 retinas). (**h**) Quantification of the percentage of tdTomato+ cells co-labeling with amacrine cell markers Pax6 and TFAP2, retinal ganglion cell marker RBPMS, and bipolar cell maker Chx10. **(i)** Percentages of tdTomato+ cells co-labeling with GFP or calretinin. **(j)** Percentage of GFP+ cells co-labeling with calretinin. n=4 measurements, 3 animals for each experiment in **h-j**.

Both the S3- and S5-stratifying Gbx2+ populations co-localized with pan-amacrine cell markers TFAP2 and Pax6 (Fig. 1d, h, Fig 1- supplement 2a), and lacked expression of the retinal ganglion cell marker RBPMS and bipolar cell marker Chx10 (Fig. 1h, Fig 1- supplement 2b, c). GFP expression could be detected in 55.3% of the total Gbx2/tdTomato+ population (Fig. 1e, i). A second marker, calretinin, labeled 53.6% of the total Gbx2+ ACs (Fig. 1f, i) and nearly all (94.9%) of the GFP+ Gbx2+ ACs were co-labeled by Calretinin (Fig. 1j). Both GFP and calretinin label the S3 sublamina of the IPL, whereas S5 is negative for both GFP and calretinin (Fig. 1e-g). Many retinal neuron subtypes form regularly spaced mosaics to provide even coverage across the visual field (Keeley *et al.*, 2020). Both S3- and S5-stratifying Gbx2+ populations formed regular mosaics in the INL and GCL independent of each other (Fig. 1 – supplement 3a-k). The GFP+/Calretinin+ S3-stratifying subtype was present at a slightly higher density than the GFP-/Calretinin-S5-stratifying subtype (Fig. 1 – supplement 3i). The cell density of both subtypes was consistent across the different areas of the retina (Fig. 1 – supplement 3l-n). These data suggest that *Gbx2*^*CreERT2-IRES-GFP*^ labels two distinct AC subtypes: a GFP+/Calretinin+ population that stratifies in S3 and a GFP-/Calretinin-population that stratifies in S5 of the IPL.

### S3- and S5-Gbx2+ amacrine cells have distinct molecular profiles

The transcriptomic profiles of related neuronal subtypes can provide clues into the molecular basis of the morphology, development, and function, as well as identify markers for distinguishing individual subtypes. To identify the molecular differences between S3- and S5-Gbx2+ ACs, we performed bulk RNA-seq on Gbx2+ ACs isolated from P8 retinas of *Gbx2*^*CreERT2-IRES-EGFP*^; *R26*^*LSL-tdTomato*^ mice administered 25µg tamoxifen at P1. We used fluorescence activated cell sorting (FACS) to separate the S3-Gbx2+ AC (GFP+/tdTomato+) and S5-Gbx2+ AC (GFP-/tdTomato+) populations (Fig. 2 – supplement 1). Bulk RNAseq was preformedon these isolated populations and identified 18 and 67 differentially expressed genes (DEGs) enriched in S3- and S5-Gbx2+ ACs, respectively (Fig. 2a, Table 1-2). *Gbx2* was expressed 13-fold higher in the S3-Gbx2+ ACs versus the S5-Gbx2+ ACs (Fig. 2b). Consistent with this result, *GFP* (driven from the endogenous *Gbx2* locus) was expressed at 10-fold higher in the S3-Gbx2+ ACs compared to the S5 subtype (Fig. 2b). Calretinin, which distinguishes S3- and S5-Gbx2+ ACs by immunohistochemistry (Fig. 1f-j), was expressed 19-fold higher in S3-Gbx2+ ACs (Fig. 2b).

**Figure 2.**
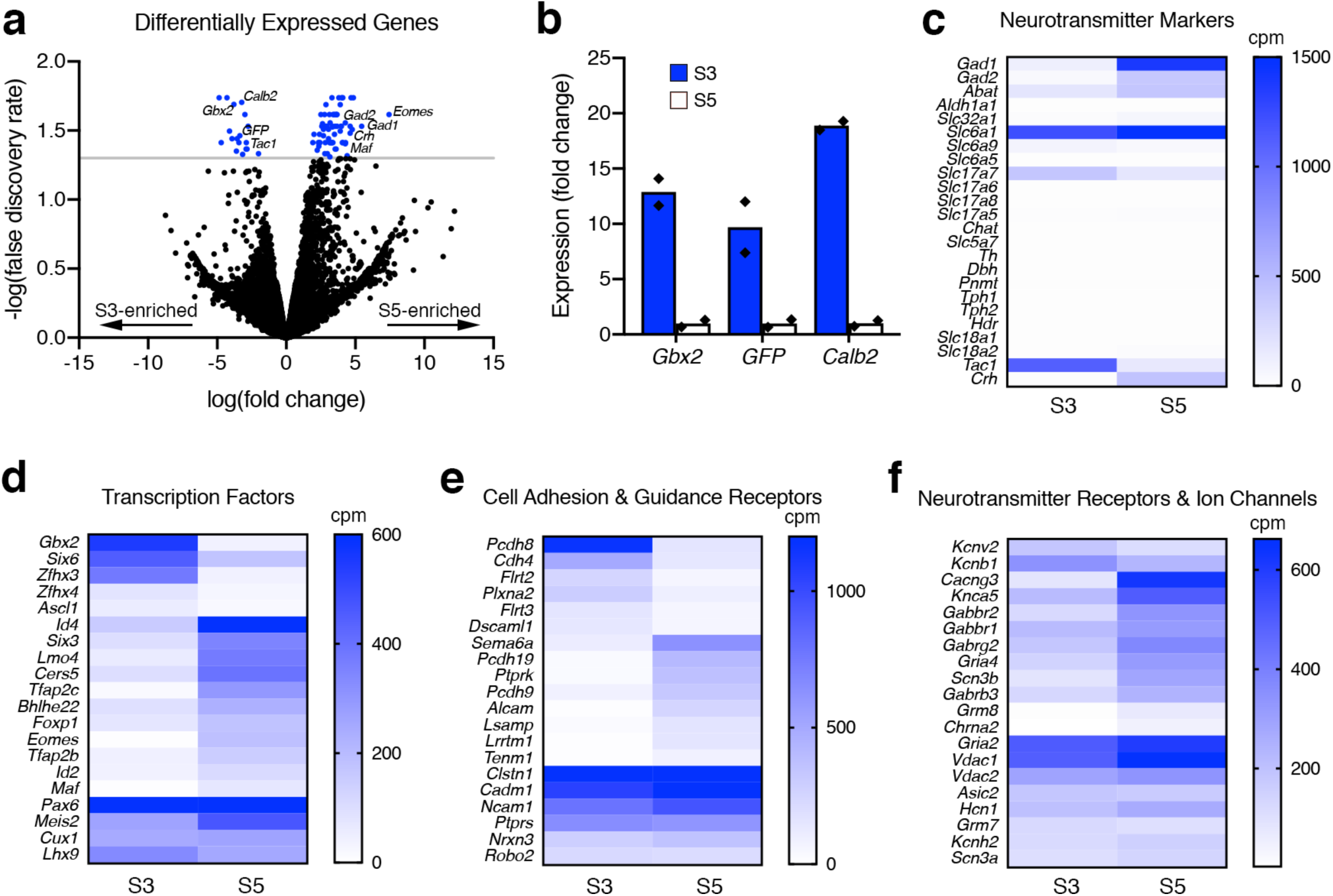
Transcriptomic profiling of S3- and S5-stratifying Gbx2+ AC subtypes identifies molecular differences. **(a)** Differentially expressed genes (DEGs) identified by RNAseq between S3- and S5-Gbx2+ ACs are displayed in a volcano plot (negative fold change, S3-enriched; positive fold change, S5-enriched). Blue points represent genes that have significant differences in between the S3- and S5-Gbx2+ AC subtypes (p<0.05, false discovery rate (FDR), gray line). **(b)** Fold-change in expression determined by RNAseq for known S3-specific markers in S3 (blue bars) and S5 (white bars) Gbx2+ AC subtypes. Each point (black diamond) represents a single sample/animal. **(c)** The mean expression of neurotransmitter markers from RNAseq between S3- and S5-Gbx2+ ACs displayed in a heatmap. Neurotransmitter markers are subdivided into GABA synthesis (*Gad1, Gad2, Abat, Aldh1a1*), GABA transport (*Slc32a1, Slc6a1*), Glycine transport (*Slc6a9, Slc6a5*), Glutamate transport (*Slc17a7, Slc17a6, Slc17a8, Slc17a5*), Cholinergic synthesis and transport (*Chat, Slc5a7*), Catecholamine synthesis (*Th, Dbh, Pnmt*), Seratonin synthesis (*Tph1, Tph2*), Histamine synthesis (*Hdc*), Monoamine transport (*Slc18a1, Slc18a2*), and Neuropeptides (*Tac1, Crh*). **(d-e)** Heatmaps comparing the mean expression of highly and differentially expressed **(d)** transcription factors, **(e)** cell adhesion and axon guidance receptors, and **(f)** neurotransmitter receptors and ion channels. Cpm, counts per million reads.

The majority of AC subtypes are categorized by their neurotransmitter profile as either GABAergic or glycinergic (Kay *et al.*, 2011). Based on the RNAseq results, the GABAergic enzymes *Gad1* and *Gad2* are highly expressed in S5-Gbx2+ ACs (Fig. 2c). In contrast, S3-Gbx2+ ACs were negative for all the standard neurotransmitter markers, including the GABAergic markers *Gad1, Gad2*, and the glycinergic markers *Slc6a9* and *Slc6a5* (Fig. 2c). Both S3 and S5 subtypes expressed low levels of the glutamatergic marker, *Slc17a7*, however this is likely due to photoreceptor contamination. The neuropeptides *Tachykinin1* (*Tac1*) and *Corticotropin-releasing hormone* (*Crh*) were highly expressed in the S3- and S5-Gbx2+ subtypes, respectively (Fig. 2c).

To gain additional insight into the differences between S3- and S5-Gbx2+ ACs, we expanded our analysis to include transcription factors (Fig. 2d), cell adhesion and axon guidance receptors (Fig. 2e), and neurotransmitter receptors and ion channels (Fig. 2f). The S3-Gbx2+ AC subtype expressed the transcription factors *Gbx2* and *Zfhx3* significantly higher than in the S5 subtype, whereas the S5 subtype selectively expressed *Id4, Lmo4, Tfap2c, Eomes*, and *Maf* (Fig. 2d). Both S3 and S5-Gbx2+ AC subtypes expressed high levels of the transcription factors *Pax6, Meis2, Cux1*, and *Lhx9* (Fig. 2d). The cell adhesion molecule *Pcdh8* was expressed significantly higher in the S3-Gbx2+ ACs, whereas *Pcdh19, Alcam, Lrrtm1*, and *Tenm1* were higher in the S5 subtype (Fig. 2e). Both shared high expression of the adhesion receptor genes *Clstn1, Cadm1, Ncam1, Ptprs, Nrxn3*, and *Robo2* (Fig. 2e). Unlike transcription factors and adhesion receptors, neurotransmitter receptors and ion channels showed little difference in expression between the two subtypes, with the exception of *Cacng3, Grm8*, and *Chrna2* genes showing significant enrichment in the S5 subtype (Fig. 2f). These differentially expressed genes identify markers to distinguish the two Gbx2+ AC subtypes, and provide insight into the function of these cells, particularly with respect to neurotransmitter release.

### S3-Gbx2+ amacrine cells are non-GABAergic, non-Glycinergic (nGnG)

The majority of ACs (∼85%) release either GABA or glycine to inhibit downstream postsynaptic neurons (Kay et al. 2011). Based on the RNAseq results, S5-Gbx2+ ACs express both *Gad1* and *Gad2* and are therefore GABAergic (Fig. 2c). We confirmed this with immunohistochemistry, which showed that 28.4% of Gbx2+ ACs in the INL co-labeled with the GABAergic marker GAD67, while the remaining Gbx2+ ACs were negative for GAD67 (Fig. 3a-c, h). The GAD67+/Gbx2+ ACs are likely S5-Gbx2+ ACs, as ∼37% of all Gbx2+ ACs in the INL are the S5 subtype (Calretinin-/GFP-) (Fig. 1 - supplement 3l). Neither Gbx2+ AC subtype co-localized with the glycinergic marker, GlyT1/2 (Fig. 3d-e, h). Therefore, S3-Gbx2+ ACs are a subtype of non-Glycinergic, non-GABAergic (nGnG) AC. These results are consistent with previous scRNAseq analysis that showed a cluster of ACs identified by their expression of Gbx2 lacked expression of *Slc6a9* or *Gad1/2* (Macosko *et al.*, 2015; Yan *et al.*, 2020). When we examined other neurotransmitter markers known to be expressed in specific AC subtypes, we found little (<1%) or no co-localization with Gbx2+ ACs (Fig. 3h, Fig. 3 - supplement 1a-d), consistent with our RNAseq results (Fig. 2c).

**Figure 3.**
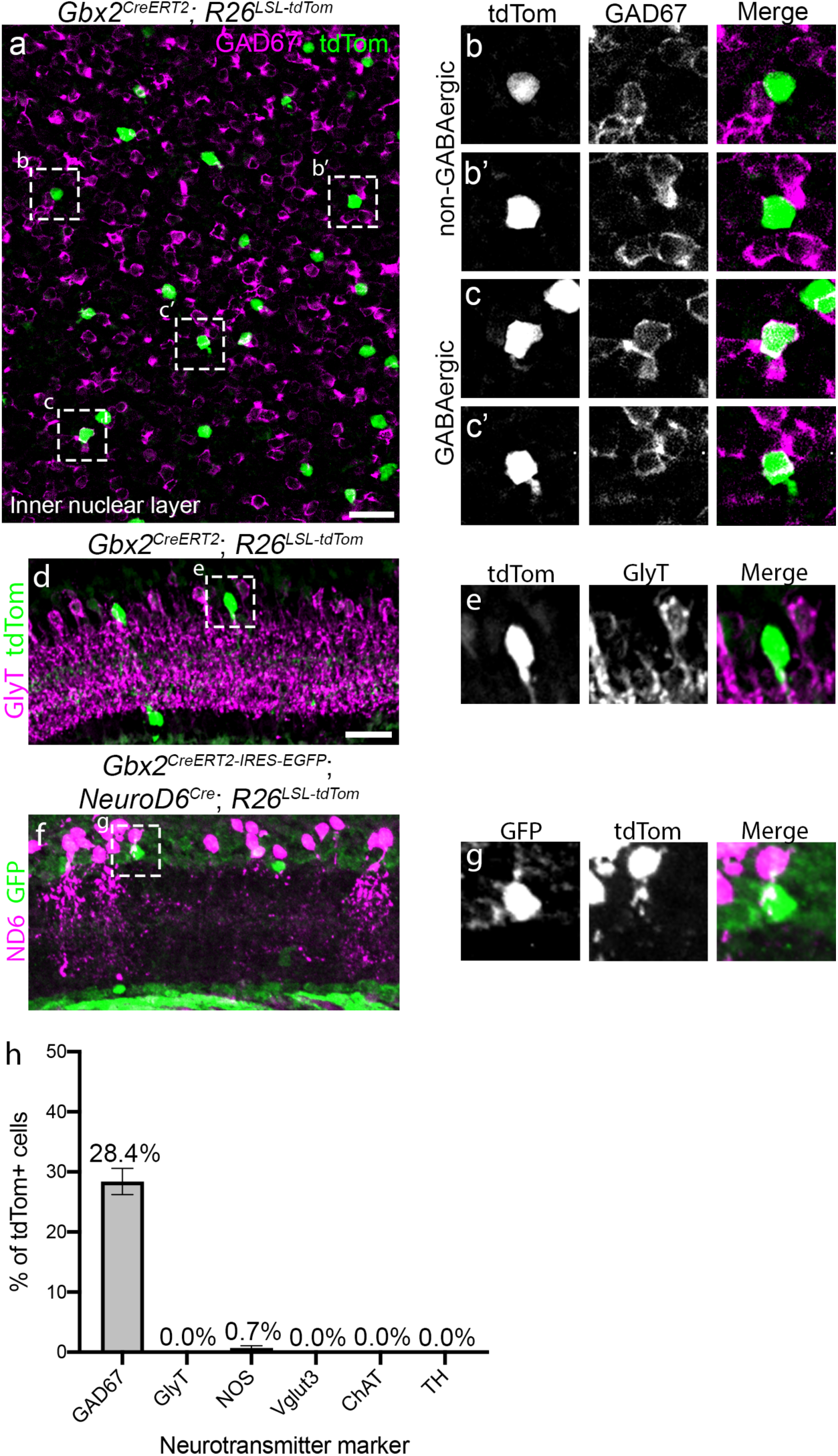
S3-Gbx2+ neurons are a subtype of non-GABAergic non-Glycinergic amacrine cell. **(a)** Inner nuclear layer of an adult retina *en face* from a *Gbx2*^*CreERT2*^; *R26*^*LSL-tdTom*^ mouse immunolabeled for tdTomato (green) and GAD67 (magenta). **(b-c)** Magnified images of Gbx2+ ACs from the panel (**a**) that do not co-localize with GAD67**(b-b’)** and GABAergic Gbx2+ ACs that colocalize with GAD67 **(c-c’). (d-e)** A retinal cross-section from a *Gbx2*^*CreERT2*^; *R26*^*LSL-tdTom*^ mouse shows no co-localization between Gbx2+ ACs with GlyT1/2. **(f)** Gbx2+ ACs (GFP, green) do not co-localize with NeuroD6+ nGnG ACs (tdTom, magenta) in retinal sections from a *Gbx2*^*CreER-ires-EGFP*^; *NeuroD6*^*Cre*^; *Rosa26*^*LSL-tdTomto*^ mouse. **(g)** Magnified image from panel (f) labeled with GFP (*Gbx2*, left), tdTomato (*NeuroD6*, middle), and a merge image (right). **(h)** Quantification of the percentage of tdTomato+ cells that co-localize with neurotransmitter markers. GAD67 quantification was performed in INL only. n > 125 neurons and 3 mice for each condition. Scale bar, 25μm in (**a, d, f**), 10μm in (**b, c, e, g**).

Previous studies have hinted at the presence of at least two nGnG AC subtypes (Cherry *et al.*, 2009; Macosko *et al.*, 2015; Yan *et al.*, 2020). One nGnG AC subtype is defined by expression of the transcription factor *NeuroD6* (Kay *et al.*, 2011). *NeuroD6*+ nGnG ACs have a narrow dendritic arbor that stratifies diffusely through multiple layers of the IPL, in contrast to the dendritic arbors of S3-Gbx2+ ACs, which have a medium sized arbor that is mono-stratified. To confirm that *Gbx2* and *NeuroD6* label distinct populations of nGnG ACs, we generated a *NeuroD6*^*Cre*^; *Gbx2*^*CreERT2-IRES-EGFP*^; *R26*^*LSL-tdTomato*^ mice and found no overlap between the NeuroD6+ (tdTomato+) and Gbx2+ (GFP+) AC subtypes (Fig. 3f-g). Therefore, S3-Gbx2+ ACs are a novel, uncharacterized subtype of nGnG AC.

### S3- and S5-Gbx2 ACs have distinct morphological properties

The dendritic morphology of a neuronal subtype is directly related to its function (Lefebvre *et al.*, 2015). Dendritic shape and size are particularly important for the computations performed by AC subtypes, as dendrites contain both pre- and post-synaptic sites. To define the morphological properties of S3- and S5-stratifying Gbx2+ ACs we used two different sparse-labeling techniques: a genetic approach using low dose tamoxifen (0.05mg), or single-cell fills with Neurobiotin (Fig. 4a-d). S3-Gbx2+ ACs had an average dendritic arbor area of 10,331±629 µm^2^ and total dendrite length of 1766±109 µm. S5-Gbx2+ ACs had significantly larger dendritic arbors, with an area of 44,643±11,699 µm^2^ and length of 2800±401 µm (Fig. 4e, Fig. 4 – supplement 3a). S3-Gbx2 ACs had a significantly higher average dendrite branch density of 117.8±16.9 branches/100 µm^2^, compared to 34.3±12.9 in S5-Gbx2+ ACs (Fig. 4g). We did not observe any difference between S3- and S5-Gbx2+ ACs in dendrite coverage factor (Fig. 4 - supplement 1d), total number of branches (Fig. 4 - supplement 1b), or branch crossovers (Fig. 4 – supplement 1c).

**Figure 4.**
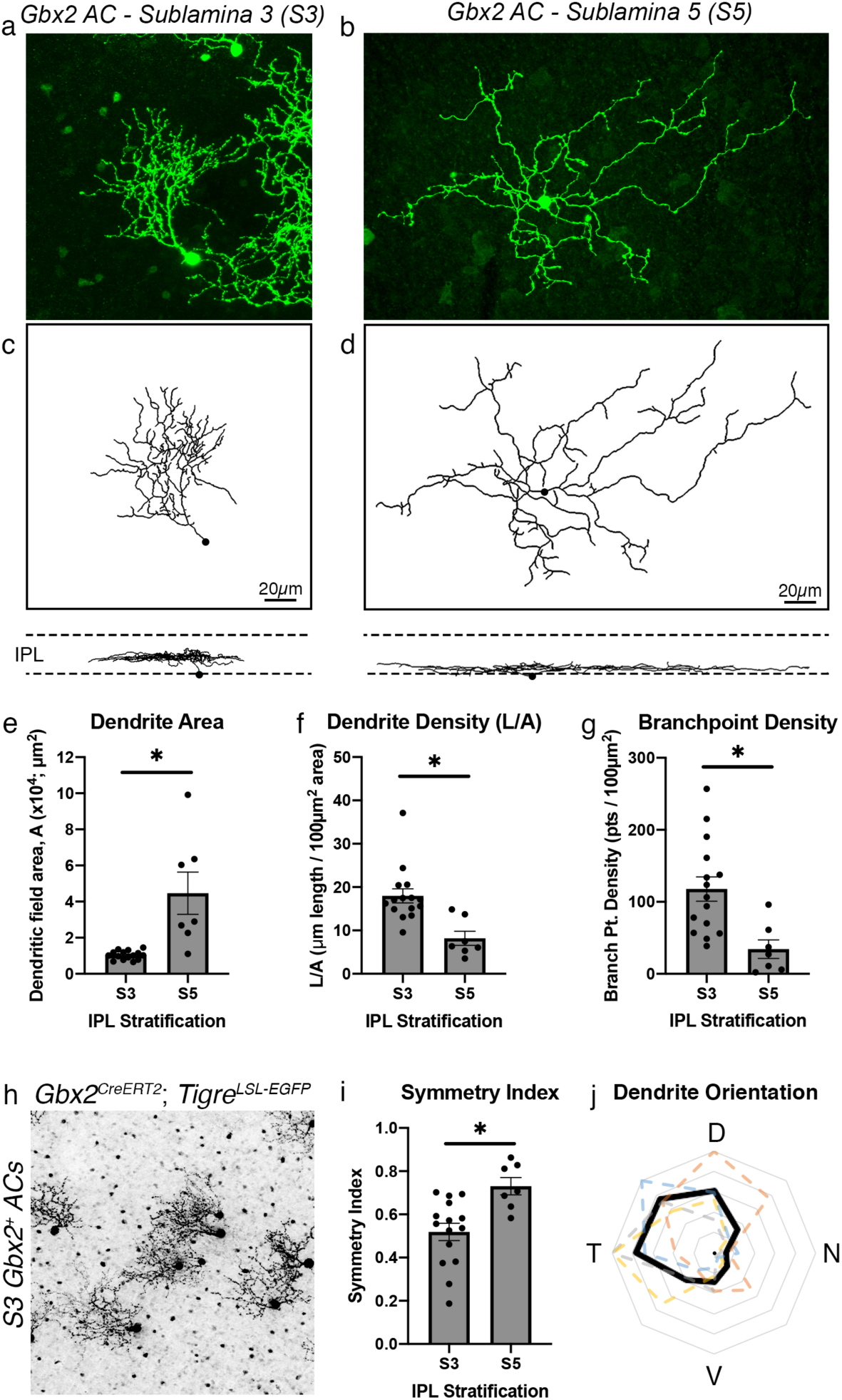
S3- and S5-Gbx2+ ACs distinct morphological features. (**a-b**) Single cell labeling of a (**a**) S3 stratifying and (**b**) S5 stratifying Gbx2+ ACs using a *Gbx2*^*CreERT2-IRES-EGFP*^; *TIGRE*^*TIT2L-GFP-ICL-tTA2*^ mouse and neurobiotin-cell fill. (**c-d**) Dendrite morphology traces constructed using the Filaments plugin in Imaris software for the Gbx2+ ACs in (**a**) and (**b**) respectively. Below are the traces in the orthogonal view to display the stratifications of each neuron in the IPL. (**e-g**) S3- and S5-Gbx2+ AC morphology in (**e**) dendrite area (p=0.0003), (**f**) dendrite density (p=0.0015), (**g**) branchpoint density (p=0.0009). Morphological data was collected from 15 S3-Gbx2+ ACs (3 animals) and 7 S5-Gbx2+ ACs (4 animals). (**h**) Gbx2+ ACs sparsely labeled in a flat-mounted P21 retina in a *Gbx2*^*CreERT2-IRES-EGFP*^; *TIGRE*^*TIT2L-GFP-ICL-tTA2*^ mouse. Quantification of dendritic arbor symmetry between S3- and S5-stratifying ACs (p=0.0039). (**j**) A polar plot of dendrite orientation of S3-targeting Gbx2+ ACs; black trace represents the mean and colored dashed traces represent neurons quantified from a single retina (n >20 neurons per retina, n= 4 retinas). The concentric gray rings represent the average percentage of neurons with dendrites extending in the specific orientation (outer most ring = 100%, 20% increments). *p<0.05 by an unpaired t-test with Welch’s correction.

Sparse labeling revealed that the dendritic arbors of S3-Gbx2+ ACs exhibited a clear asymmetry, with the arbor rarely exceeding 180 degrees (Fig. 4h). As a population, S3-Gbx2+ ACs had a significantly lower symmetry index than S5-Gbx2+ ACs (Fig. 4i). We analyzed the dendrite orientation of S3-Gbx2+ ACs at a population level and found that their asymmetric dendrites were consistently oriented in a dorsal-temporal direction (Fig. 4j). Both dendrite asymmetry and orientation of S3-Gbx2+ ACs were consistent across all quadrants of the retina (Fig. 4 – supplement 1e-f, 1g-j, respectively). In contrast to S3-Gbx2+ ACs, the S5-Gbx2+ ACs did not show any obvious asymmetry or orientation bias (Fig. 4i). Overall, these data show that S3- and S5-Gbx2+ ACs exhibit significantly different morphological features, supporting the hypothesis that they are distinct subtypes.

### S3-Gbx2+ amacrine cells are tracer coupled to bipolar cells

Many amacrine cells show selective electrical coupling to other neurons, including bipolar cells, ganglion cells, and other amacrine cells (Bloomfield & Volgyi, 2009). Electrical synapses play a major role in the transmission of visual information in the retina(O’Brien & Bloomfield, 2018), and are typically revealed by filling cells with Neurobiotin, a small tracer molecule that is permeable through most gap-junctions (Vaney, 1991). When we filled S3-Gbx2+ AC somas in the ganglion cell layer with Neurobiotin we found that the tracer spread into a sparse number of neighboring retinal neurons with somas in the INL (Fig. 5a-b). The coupled cells could be identified as bipolar cells based on their morphology in orthogonal optical sections (Fig. 5g). At higher magnification we also observed tracer-filled bipolar cell axon terminals overlapping S3-Gbx2+ AC dendrites in the same optical section *en face* (Fig. 5h). In contrast to the S3-Gbx2 ACs, S5-Gbx2+ ACs did not show consistent tracer coupling with other retinal cells (Fig. 5j).

**Figure 5.**
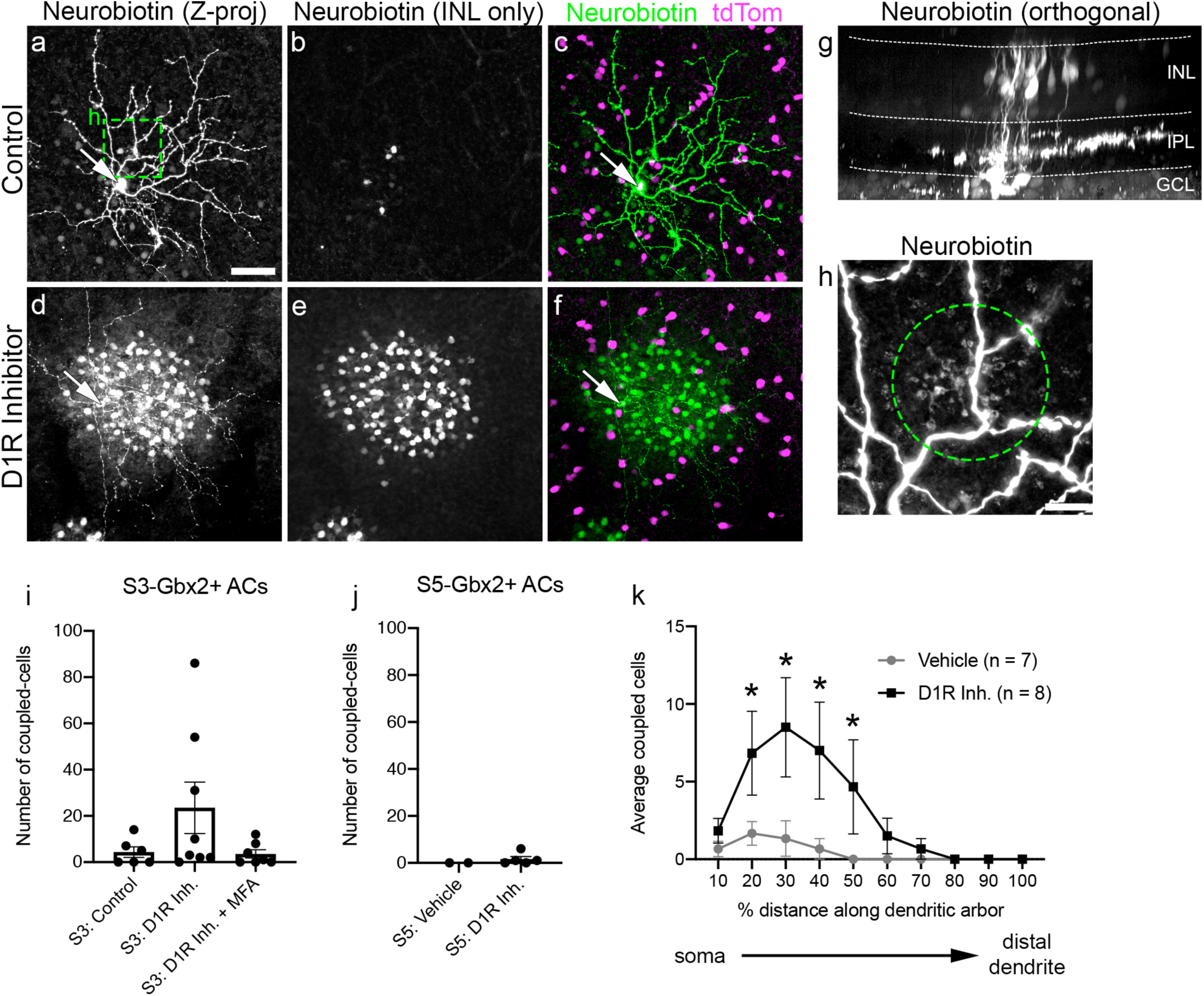
Gbx2+ ACs are electrically coupled to neighboring bipolar cells. Neurobiotin-filled S3-stratifying Gbx2+ AC (arrows, cell bodies) incubated in (**a-c**) vehicle and (**d-f**) D1 receptor inhibitor, 50µM SCH23390, in flat-mounted retinas. (**a**,**d**) Z-projection through the entire retina. (**b**,**e**) Z-projection through INL only. (**c**,**f**) Merged Z-projection with neurobiotin (green) and Gbx2+ ACs (magenta). **(g)** Orthogonal view of neurobiotin filled Gbx2+ AC (arrow) and dye-coupled bipolar cells (BCs). **(h)** Magnified image (box in a) of BC axon terminals (circle) co-localizing with a dendritic branch of neurobiotin-filled Gbx2+ AC. **(i-j)** Mean number neurons dye-coupled to a single (**i**) S3- and (**j**) S5-stratifying Gbx2+ AC (S3-Vehicle, n= 6 cell-fills; S3-D1R Inh., n=8; S3-D1R Inh. + MFA, n=7; S5-Vehicle, n=2; S5-D1R Inh., n=3; Welch’s t-test). **(k)** The spatial distribution of dye-coupled cells to a single S3-stratifying Gbx2+ AC along its dendritic arbor. *p<0.05 calculated by a Two-way ANOVA with multiple comparisons. Scale bar, 50μm in **a-g**, 10μm in **h**.

The extent and strength of coupling in the retina is linked to the ambient light level through the release of the neuromodulator dopamine. Dopamine release increases as the background light-level increases (Hampson *et al.*, 1992). The increased concentration of dopamine activates D1 receptors in all AII ACs, reducing their coupling to BCs. To determine whether S3-Gbx2+ AC to bipolar cell coupling is also modulated by dopamine, we measured coupling in the presence of the D1 receptor antagonist, SCH23390 (50µM). Inhibition of D1 receptors dramatically increased coupling to bipolar cells in 3 out of 8 S3-Gbx2+ ACs, while coupling in the remaining S3-Gbx2+ ACs was unaffected (Fig. 5d-i). The specificity of tracer coupling was confirmed by bath application of the gap junction antagonist Meclofenamic acid (100µM MFA), which blocked coupling in both baseline and D1 receptor antagonist conditions (Fig. 5i).

The coupling between S3-Gbx2+ ACs and bipolar cells show unusual spatial features. In other ACs, coupling typically occurs homotypically and spreads laterally to serve a signal averaging function (Bloomfield & Volgyi, 2009). In contrast, we never observed S3-Gbx2+ ACs coupled to neighboring S3-Gbx2+ ACs, and their coupling to BCs appeared to be spatially selective. BCs coupled to S3-Gbx2+ ACs were clustered towards the center of the dendritic arbor (Fig. 5k). Even with addition of the D1R antagonist, bipolar cell coupling to Gbx2+ ACs never extended into the distal 20% of the dendrite arbor (Fig. 5j). Inhibition of the D1 receptors did not unmask coupling between the S5-Gbx2+ ACs and other retinal neurons (Fig. 5j).

### Input differences between S3- and S5-Gbx2+ ACs

We next sought to characterize the intrinsic physiological properties of S3- and S5-Gbx2+ ACs. S3-Gbx2+ ACs cells stratify at the border between the On and Off sublaminae of the IPL and thus potentially receive input from both On- and Off-type BCs. The S5-Gbx2+ ACs cells stratify within the On-sublamina, and are therefore likely to receive input from On-type bipolar cells. To test these predictions we recorded light evoked postsynaptic potentials (PSPs) from S3- and S5-Gbx2+ ACs. Cells were stimulated with light spots of 50 to 80% contrast, square-wave modulated at 2Hz, centered on the receptive field. The background intensity was set at a photopic level (10^3^ photons/µm^2^/s). Consistent with the stratification levels of their dendrites, S3-Gbx2+ ACs cells showed strong depolarization during both the On phase (increase in luminance) and Off phase (decrease in luminance) of the stimulus (black trace, Fig. 6a, left). In contrast, S5-Gbx2+ ACs cells were depolarized only during the On phase, and are hyperpolarized during the Off phase (black trace, Fig. 6a, right). Thus, the physiological inputs are consistent with the stratification level of the dendrites for both S3- and S5-Gbx2+ ACs.

**Figure 6.**
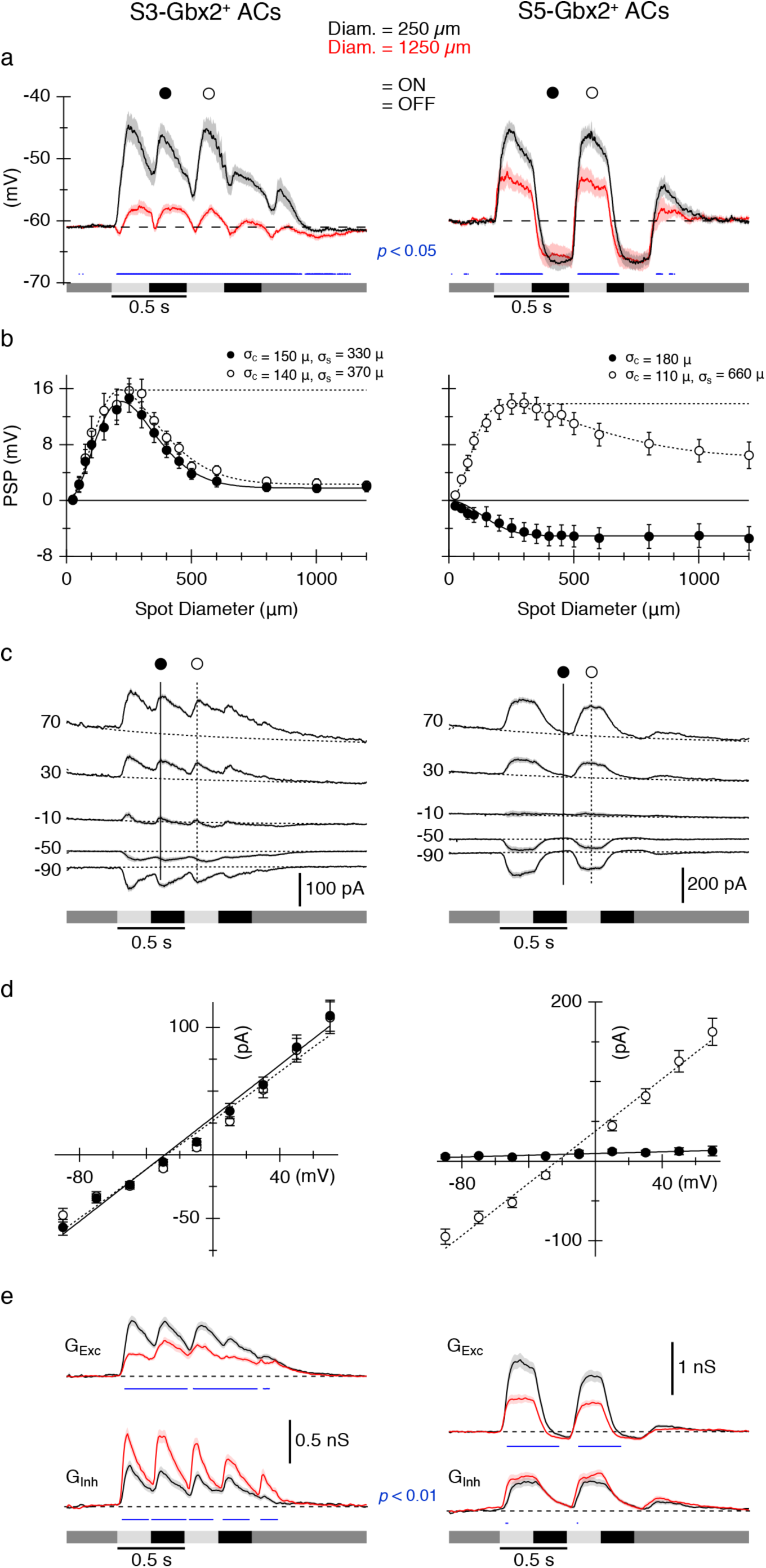
S3- and S5-Gbx2+ ACs exhibit distinct spatial receptive field properties and On-Off inputs. **(a)**.Average light-evoked postsynaptic potentials (PSPs) in S3 (n=12) (**left**) and S5 (n=17) (**right**) Gbx2+ ACs in response to a 250 *μ*m (black) or 1200 *μ*m (red) diameter spot stimulus. Luminance indicated by shaded bars underneath traces. (**b)** Area response functions from S3 (**left**) and S5 (**right**) cells measured as the amplitude of the PSPs versus stimulus diameter at indicated time points in **(a)**. Surround suppression of PSPs was 86% for S3 and 53% for S5 cells. **(c)** Average light-evoked postsynaptic currents (PSCs) in (n=56) (**left**) S3 and (n=45) (**right**) S5 cells for center spot stimuli during a series of voltage steps from -90 to +70mV. **(d)** I-V relations for the PSCs at the time points indicated in ***(c).*** The dotted and solid lines show average least-squares fits to the I-V relations. **(e)** Average synaptic conductance traces calculated from fits to I-V relations measured every 10 ms during the light stimulus (see Methods). Traces and symbols represent average responses. Shading and error bars represent SEM. Blue traces in **(a)** and **(e)** indicate significant differences (paired t-test) between the solid and dashed data. P values indicated on panels.

### S3- and S5-Gbx2+ ACs cells receive center and surround inhibition

In many retinal neurons the spatial structure of the receptive field is shaped by center/surround organization, in which illumination outside a neuron’s excitatory center receptive field elicits an antagonistic reponse. We presented spots with a range of diameters to test for differences between the spatial extent of the two cell types’ receptive fields. The largest spots suppressed center responses in both cell types. For S3-Gbx2+ ACs, maximal surround stimulation suppressed On and Off PSPs equally by 86% (Fig. 6a-b, left), whereas surround suppression was weaker in the S5-Gbx2+ ACs, reducing center PSPs by about 53% (Fig. 6a-b, right). Activation of the surround had little effect on the amplitude of the hyperpolarization evoked during the OFF-phase of the stimulus (Fig. 6b, left, filled symbols). The spatial extent of the center and surround were estimated at fixed time-points during the On and Off phase of the stimulus by fitting a difference of Gaussians (DOG) function to the amplitude of the responses. The On and Off responses of the S3 ACs showed essentially identical spatial tuning profiles, with the extent of the surround being about 2.5-fold wider than the center (Fig. 6b, left). Maximum dendritic diameter averaged 181 ± 8 µm (n=17), as estimated from morphological analysis, which compares well with the physiological center size of ∼145µm, estimated as the space constant from the DOG fit. The S5 ACs were more divergent. The maximum dendritic diameter averaged 258 ± 12 µm (n=27), while the physiological center size estimated during the On-phase was only ∼110 µm (Fig. 6b, right), and was smaller than the center size measured during the Off-phase (∼180µm). A similar difference is apparent in a previous study (Park et al., 2018).

To examine potential synaptic mechanisms providing surround antagonism, we recorded light evoked postsynaptic currents (PSCs) in response to center-only and full-field visual stimuli, and calculated the component excitatory and inhibitory inputs (Fig 6c-e). Excitation and inhibition were activated during the On and Off phase of the stimulus for S3-Gbx2+ ACs, and appeared to have very similar temporal dynamics (Fig. 6e, left). The S5 ACs showed a similar pattern, with excitation and inhibition showing similar temporal wave-forms. However, while inhibition was slow to turn-off during the Off-phase, excitation declined more rapidly and dipped below zero (Fig. 6e, right), presumably driving the hyperpolarization seen during the Off-phase in Fig. 6a. Comparison of the center-only and full-field responses provides insight into the mechanisms of surround antagonism. For the S3 ACs, excitation was suppressed and inhibition enhanced for full-field stimuli relative center-only stimuli (Fig. 6e, red vs black, left), suggesting that the antagonistic surround shown in Fig. 6a results from complementary changes in both excitation and inhibition. For S5 ACs, by contrast, activation of the surround suppressed only the excitation, and had no significant effect on the inhibitory input (Fig. 6e, red vs black, right).

### S3- and S5-Gbx2+ ACs receive distinct On and Off pathway excitation and inhibition

These data suggest excitation and inhibition may arise through both the On and Off pathway for S3-Gbx2+ ACs, but only the On pathway for S5-Gbx2+ ACs. Consistant with this hypothesis, when the On-pathway signaling was blocked by using L-AP4 to block photoreceptor inputs to On-bipolar cells, light-evoked synaptic input to S5-Gbx2+ ACs was abolished (Fig. 7a, right). In contrast, the light-evoked synaptic inputs to S3-Gbx2+ ACs during the On-phase of the stimulus were strongly suppressed, while the Off pathway excitation and inhibition were largely unaffected. (Fig. 7a, left). Inhibition is in-phase with excitation during On and Off responses in S3-Gbx2+ ACs and during On responses in S5-Gbx2+ ACs suggesting that it could serve to regulate the gain of signaling by counterbalancing excitation.

**Figure 7.**
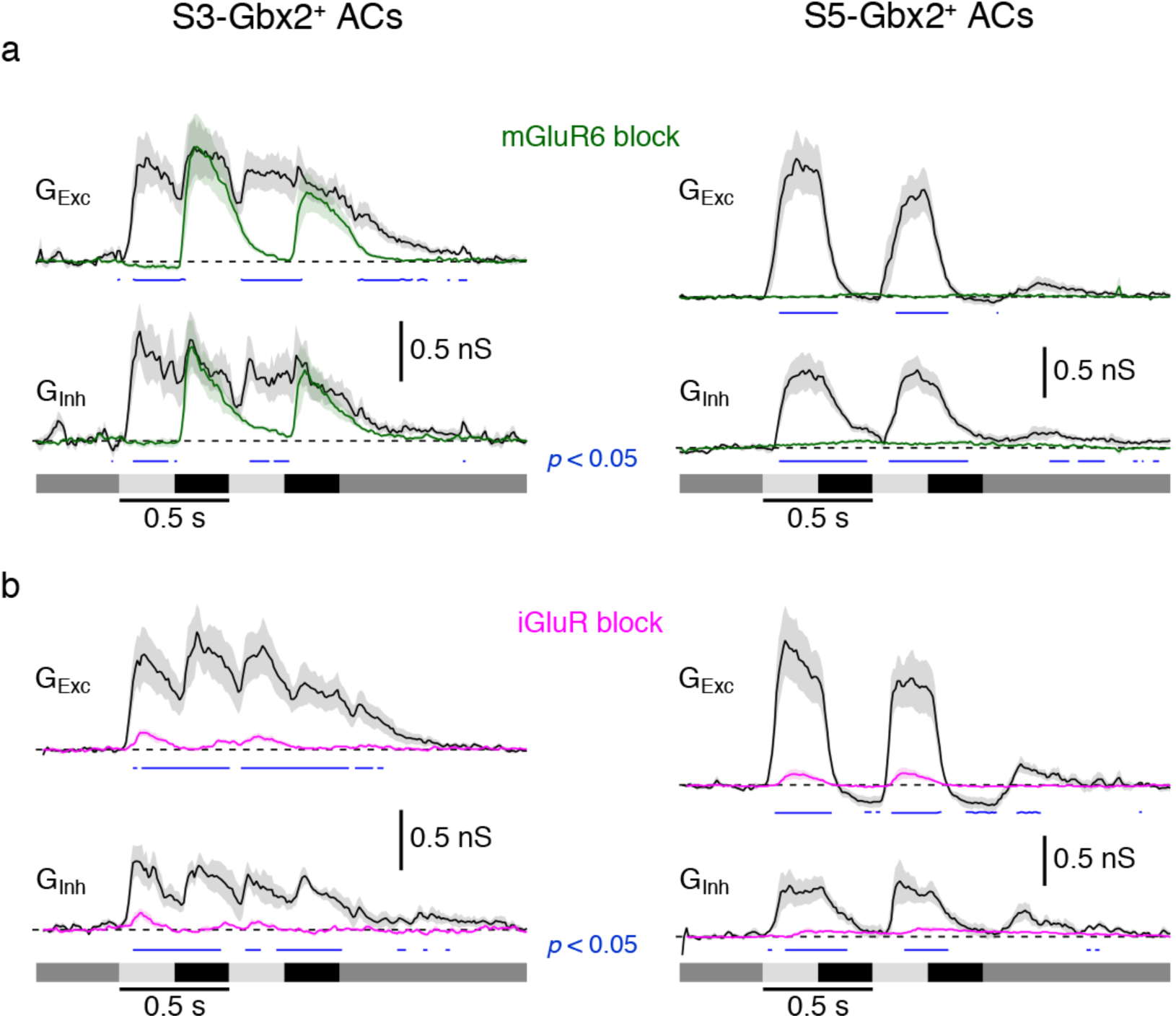
S3- and S5-Gbx2+ AC subtypes have distinct excitatory and inhibitory inputs. **(A)** Conductance measurements from (n=7) S3 (**left**) and S5 (**right**) cells in response to a center spot stimulus in the presence (green) or absence (black) of mGluR6 receptor agonist 10 μM L-AP4. **(B)** Conductance measurements from (n=6) S3 and S5 cells in response to a center spot stimulus in the presence (magenta) or absence (black) of a cocktail of iGluR receptor agonists: 1 μM ACET, 20 μM GYKI, and 50 μM D-AP5. Traces represent average responses. Shading represents SEM. Blue traces indicate significant differences (paired t-test) between the drug and control conditions. P values indicated on graphs (p< 0.05).

The S3- and S5-Gbx2+ ACs show distinct Neurobiotin-coupling patterns to peri-somatic bipolar cells (Fig. 5). To test whether these gap junction connections support significant, light-evoked electrical synaptic input to the ACs, we blocked glutamatergic inputs using a cocktail of antagonists, including the NMDA receptor antagonist D-AP5 (50μm), and the AMPA and Kainate receptor antagonists ACET and GYKI (1μM and 50μM). Under these conditions, mGluR6 mediated transmission to On-type bipolar cells is preserved, and if gap junctions mediate electrical transmission to S3- or S5-Gbx2+ ACs, excitatory On-responses in these cells should also be preserved. However, light-evoked responses using stimulus spots large enough to activate all coupled bipolar cells are almost completely blocked (Fig. 7b, magenta traces), suggesting that excitatory drive through any electrical synapses from the coupled BCs is functionally insignificant under these recording conditions. These experiments don’t rule out the possibile presence of functionally significant electrical synapses between Off-bipolar cells and S3-Gbx2+ ACs, because AMPA/Kainate antagonists block cone input to Off bipolar cells. However, current-voltage relations of excitatory synaptic inputs during light-off responses, recorded with inhibition blocked, were linear and reversed near zero millivolts, which is inconsistent with a major input through gap junctions (data not shown).

## DISCUSSION

We have identified two AC subtypes in the mouse retina genetically labeled by the *Gbx2*^*CreERT2-IRES-EGFP*^ mouse line that are morphologically, physiologically, and molecularly distinct from each other. S3-Gbx2+ ACs are On-Off type cells (Fig. 5), have smaller, asymmetric dendrites that stratify in sublamina 3 of the IPL (Fig. 4), and exhibit heterotypic gap junction coupling (Fig. 6). S5-Gbx2+ ACs are On-type cells, have larger, more symmetric dendrites that stratify in sublamina 5 of the IPL, and lack consistent gap junction coupling. S3-Gbx2+ ACs lack expression of any of the standard inhibitory or excitatory neurotransmitters, identifying them as a new subtpe of nGnG ACs, whereas the S5-Gbx2+ ACs are GABAergic. RNAseq revealed the distinct molecular profiles of these two Gbx2+ AC subtypes, which confirms their identification as separate sub-types and may inform future analyses of the cell-specific mechanisms that underlie their distinct morphology and physiological features.

### Categorization of amacrine cell subtypes

Previous studies have used Golgi staining, cell fills, and sparse genetic labeling to categorize many amacrine cell subtypes based on their morphology (MacNeil *et al.*, 1999; Badea & Nathans, 2004; Lin & Masland, 2006). Reconstruction of a 114µm x 80µm volume of mouse retina by serial-block face electron microscopy indicated the presence of >40 subtypes of amacrine cells (Helmstaedter *et al.*, 2013). However, these approaches may undersample rare neurons, and do not provide a means to prospectively identify and manipulate individual subtypes. A recent single-cell transcriptomic study provides evidence for the existence of 63 molecularly distinct subtypes, indicating there is greater subtype diversity than previously predicted (Yan *et al.*, 2020). One of the challenges going forward will be to match these molecularly-defined subtypes with their morphological counterparts and begin to tease out their connectivity patterns with visual circuits.

Single cell transcriptomic studies of AC subtypes initially identified two different nGnG AC subtypes, one that is molecularly similar to Glycinergic ACs and one that is similar to GABAergic ACs (Cherry *et al.*, 2009; Macosko *et al.*, 2015). The “Glycinergic-like” nGnG AC subtype is marked by specific expression of NeuroD6 (Kay *et al.*, 2011). A recent study identified two additional nGnG subtypes closely related to NeuroD6+ nGnG ACs (Yan *et al.*, 2020). In contrast, the S3-Gbx2+ AC subtype is the previously uncharacterized “GABAergic-like” nGnG subtype cluster #4 in the Macosko dataset and nGnG #4, cluster #36 in the Yan dataset. S3-Gbx2+ nGnG ACs appear to be conserved in primates based on cross-species analysis of single cell transcriptomics (Peng *et al.*, 2019).

S5-Gbx2+ ACs closely resemble cluster #7 from the Macosko study based on the selective expression of *Maf, Cxcl14, Id4*, and *Lmo4* in both transcriptomic datasets (Fig. 2a). S5-Gbx2+ ACs also share morphological, molecular, and physiological properties with the previously described CRH-1 AC (Zhu *et al.*, 2014; Jacoby *et al.*, 2015; Park *et al.*, 2018). S5-Gbx2+ ACs and CRH-1 ACs have dendritic arbors of a similar size and shape that stratify in S5 of the IPL, they both express the neuropeptide *Crh* (*corticotrophin-releasing hormone*), and have similar physiological properties in response to ON light stimuli (Jacoby *et al.*, 2015; Park *et al.*, 2018). The CRH-IRES-Cre line used in these studies labels at least three different AC subtypes, which can be distinguished based on their morphological differences. An intersectional approach pairing the *Gbx2*^*CreERT2-IRES-EGFP*^ with a recently developed *Crh-IRES-FlpO* line (Jackson Labs strain #031559) and a dual-recombinase reporter line should selectively label S5-Gbx2+ ACs and allow the direct testing of whether they are the same subtype as CRH-1 ACs.

### Dendrite morphology and Gbx2+ AC function

The receptive field properties of ACs are dependent on the morphologies of their dendritic arbors. Our electrophysiological recordings show that the morphologies and the spatial receptive field properties of the S3- and S5-Gbx2+ ACs are quite different from one another (Fig. 6a-b). Although both cells display a center-surround organization, with fairly similar center diameters, the extent of the surround is much smaller for the S3 than the S5 cells. For S3-Gbx2+ ACs, responses during both the On and Off phase of the stimulus were strongly suppressed by the surround, with space constants around ∼350 µm (Fig. 6b). The S5-Gbx2+ ACs showed much weaker surround suppression, with a space constant exceeding 650µm. With stronger surround suppression, activated with a shorter space-constant, the S3 cells will be tuned to higher spatial frequencies than the S5 cells. It seems likely that the narrow surround inhibition of the S3 cells arises in the IPL, while the broader surround of the S5 ACs probably reflects, at least in part, the center-surround organization of the cone photoreceptor output, generated by horizontal cell feedback in the OPL. The lack of any surround suppression during the Off-phase of the response (Fig. 6b) is expected if the response were produced by turning off presynaptic On bipolar cells that provide a tonic excitatory input to the S5 ACs. The disparity in the center size estimates for the On and Off phases of the stimulus might be explained by inputs from two populations of On-bipolar cells with different spatial distributions across the dendritic arbor. Further work will be required to test this hypothesis. Unlike S5 ACs, CRH-1 ACs have been shown to lack of any surround suppression during the On-phase of the response (compare Fig. 3 of (Park *et al.*, 2018), with Fig. 6a). This discrepancy probably reflects differences in the physiological conditions of the preparations rather than evidence that S5 Gbx2+ ACs and CRH-1 ACs are different subtypes.

Unlike the S5-Gbx2+ ACs, the dendritic arbors of the S3-Gbx2+ ACs are asymmetric along the dorsal-temporal axis, with the soma offset relative to the dendritic arbor. Such asymmetries can influence the response properties of neurons. For example, J-RGCs display a similar asymmetry, which appears to enhance directional responses to ventral motion (Kim *et al.*, 2008; Liu & Sanes, 2017). Examples of functional asymmetry are also found outside of the retina, such as in *Drosophila* proprioceptors, which are spatially tuned for the direction of musle movement (He *et al.*, 2019). It remains to be determined whether the asymmetric dendrites of S3-Gbx2+ ACs result in anisotropic physiological receptive field structure. The developmental mechanisms that instruct dendrite asymmetry and orientation are poorly understood, but likely involve a combination of selective directional growth and pruning of developing arbors (Liu & Sanes, 2017).

As a transcription factor, *Gbx2* is poised to regulate the cellular identity and development of Gbx2+ ACs. Subtype-specific transcription factors in retinal neurons have an important role establishing correct cell body location, dendrite morphology, and synaptic connectivity (Kay *et al.*, 2011; Whitney *et al.*, 2014; Peng *et al.*, 2017; Liu *et al.*, 2018; Peng *et al.*, 2020). *Gbx2* regulates the specification, migration, and axon guidance of several neuronal populations in the brain and spinal cord (Chen *et al.*, 2010; Luu *et al.*, 2011; Chatterjee *et al.*, 2012; Mallika *et al.*, 2015). Therefore, *Gbx2* may potentialy regulate development of Gbx2+ ACs through regulation of effector genes specific to this AC subtype, but its exact function in AC identity and development remains to be determined.

### Modes of neurotransmission in S3-Gbx2+ ACs

Without GABA, glycine, and other chemical neurotransmitters, how do S3-Gbx2+ ACs communicate with other neurons? One possibility is that these neurons are in fact GABAergic, and the very low levels of *Gad1* and *Gad2* in these cells is sufficient for low levels of GABA synthesis. There are also non-canonical pathways for GABA synthesis, although S3-Gbx2+ ACs do not express appreciable levels of *Abat* (*GABA transaminase*) or *Aldh1a1* (*aldehyde dehydrogenase 1a1*), the enzymes required for these pathways (Tritsch *et al.*, 2012; Tritsch *et al.*, 2014; Kim *et al.*, 2015). Alternatively, S3-Gbx2+ ACs could release GABA without *de novo* synthesis. Midbrain dopaminergic lack expression of *Gad1* and *Gad2* expression, yet release GABA that is taken up from the extracellular environment by plasma membrane GABA transporters (Tritsch *et al.*, 2014). S3-Gbx2+ ACs express relatively high levels of *Slc6a1*, which encodes the plasma membrane GABA transporter Gat1 (Fig. 2c).

In addition to chemical neurotransmitters, many retinal neurons also express neuromodulators and neuropeptides. Our RNA-seq results revealed that the S3-Gbx2+ ACs highly express the neuropeptide gene *Tachykinin 1 (Tac1)*, that encodes several neuropeptides, the most common one being Substance P (Fig. 2c). Release of substance P from ACs can modulate the excitability of downstream retinal ganglion cells, however at a slower time scale than most neurotransmitters (Zalutsky & Miller, 1990). The genetic identity of Substance P releasing ACs in the mouse retina is unknown, and several other AC subtypes also express *Tac1* in addition to S3-Gbx2+ ACs (Yan *et al.*, 2020). Ultimately, directly testing whether S3-Gbx2+ ACs use standard chemical neurotransmitters and/or neuromodulators will require identifying their downstream synaptic partners.

Many ACs exhibit homotypic coupling with neighboring ACs of the same subtype as seen with AII and nNOS-2 AC subtypes (Vaney, 1991; Jacoby & Schwartz, 2018). Coupling can spread signals laterally and may serve a “signal averaging” function in coupled cells. However, such signal averaging seems unlikely in S3-Gbx2+ ACs, because we never observed homotypic coupling despite significant overlap in their dendritic arbors (Fig. 7c). Rather, S3-Gbx2+ ACs are heterotypically coupled with neighboring bipolar cells (BCs). Heterotypic coupling between ACs and bipolar cells is unusual and has only been observed in glycinergic AII and A8 ACs with cone BCs (Vaney, 1991; Yadav *et al.*, 2019). The spatial pattern of the S3-Gbx2+ AC:BC tracer coupling is also unusual, as it never extends beyond the central 80% of the dendritic arbor of the S3-Gbx2+ AC. Although the physiological recordings (Figure 7b), indicate that electrical synapses contribute minimal excitatory input to S3-Gbx2+ ACs at phototopic light levels, it remains possible that signaling between S3-Gbx2+ ACs and cone BCs through electrical synapses could become prominent under specific conditions. Indeed, tracer coupling to BCs increased in a subset of S3-Gbx2+ ACs during application of a D1 receptor antagonist, which mimicks low dopamine release that occurs under scotopic conditions (Fig. 5d-f, i). These results demonstrate that tracer coupling can be modulated, and suggest that the physiological role of gap-junctions may depend on adaptation level. It is also possible that gap junction connections may mediate S3-Gbx2+ AC to excite ganglion cells indirectly by depolarizing BCs. In this situation, electrical synapses could amplify local signaling between S3-Gbx2+ ACs and BCs. The identity of the connexins that mediate coupling between S3-Gbx2+ ACs and BC remains unknown. Our RNAseq data, *Gjd2* (Cx36), *Gjc1* (Cx45), and *Gje1* (Cx23) are all expressed in S3-Gbx2+ ACs, implicate a number of potential targets.

### Potential visual modalities requiring Gbx2+ ACs

What are the functions of Gbx2+ ACs in the retina? While this remains an open question, the stratification pattern of Gbx2+ AC dendrites to specific lamina in the IPL restricts their potential synaptic partners and allows for some speculation. Each sublamina in the IPL contains the dendrites of a specific subset of RGC subtypes and responds to a specific set of visual modalities (Roska & Werblin, 2001). If S5-Gbx2 ACs are the CRH-1 subtype as hypothesized above, we would expect that they provide inhibition onto “sustained” Suppressed-by-Contrast (SbC) RGCs and ON αRGCs (also known as the M4 subtype of intrinsically-photosensitive RGCs (ipRGCs)) (Estevez *et al.*, 2012; Jacoby *et al.*, 2015; Jacoby & Schwartz, 2018; Park *et al.*, 2018). Sublamina 5 also contains the dendrites of M2, M3, and M5 subtypes of ipRGCs, which are involved in visual modalities involved in light avoidance, pupillary reflex, and circadian rhythm behaviors (Schmidt *et al.*, 2011; Sonoda *et al.*, 2020).

In contrast to the S5-Gbx2+ ACs, the S3-Gbx2+ ACs do not resemble any previously described AC subtype. With dendrites that ramify in sublamina 3 of the IPL (Fig. 1), the S3-Gbx2+ ACs could be connected to W3B RGCs that are involved in object motion detection (Zhang *et al.*, 2012; Krishnaswamy *et al.*, 2015), or other small-field RGCs involved in spatial vision (Jacoby & Schwartz, 2018). However, it is important to note that not all ACs and RGCs that co-stratify in an IPL sublamina are synaptically connected (Krishnaswamy *et al.*, 2015). Furthermore, we currently lack a complete accounting of the dendritic stratification of the 40+ RGC subtypes that have been defined molecularly, suggesting there are other potential synaptic partners (Rheaume *et al.*, 2018; Tran *et al.*, 2019). Ultimately, the presence of functional connections between S3- and S5-Gbx2+ ACs and their potential postsynaptic targets will require paired recordings, which should be feasible now that we have genetic tools to prospectively identify Gbx2+ ACs.

Over the past few years, single-cell transcriptomics has greatly increased the cellular inventory of molecularly distinct neuronal subtypes. Going forward we will need genetic tools to interrogate specific neuronal subtypes within a given neural circuit, as we have done in this study. The *Gbx2*^*CreERT2-IRES-EGFP*^ line has allowed us to define the molecular, morphological, and physiological properties of two AC subtypes. In particular, the unusual properties of the S3-Gbx2+ AC subtype raise a number of intriguing questions about their form and function within the retina for future studies.

## METHODS AND MATERIALS

### Animals and Animal Procedures

*Gbx2*^*CreERT2-IRES-EGFP*^ (Chen *et al.*, 2009), *NeuroD6*^*Cre*^ (NEX^Cre^) (Schwab *et al.*, 2000), *Ai9/Rosa26*^*LSL-tdTomato*^ (Madisen *et al.*, 2010), and *Ai140D/TIGRE*^*TRE2-LSL-GFP, CAG-LSL-tTA2*^ (Daigle *et al.*, 2018) mice were maintained on a C57BL/6J background. Tamoxifen was administered at E16, P0, and adults (>P28) ages. For E16 timepoints, 200µL of 5mg/mL tamoxifen and 2.5mg/mL progesterone dissolved in sunflower seed oil was administered to pregnant dam by oral gavage. Pregnancies were timed by the presence of a vaginal plug marked as E0.5. At P0 timepoints, 50 µL of 0.5mg/mL tamoxifen was injected with into the milk pouch of the mouse pups, as previously described (Pitulescu *et al.*, 2010). At adult ages, 200µL of 5mg/mL tamoxifen was administered by oral gavage for at least two consecutive days to ensure complete recombination of the *Cre*-dependent reporter (Fig. 1 – supplement 1). All animal procedures were approved by Oregon Health & Science University Institutional Animal Care and Use Committee, the Institutional Animal Care and Use Committee of University of California (Berkeley, CA), and conformed to the National Institutes of Health’s *Guide for the Care and Use of Laboratory Animals*.

### Immunohistochemistry

Adult retinas were prepared for immunolabeling by removing the eyes from the head of a recently euthanized mouse. The cornea was removed and the eyes were fixed in 4% EM-grade paraformaldehyde (PFA) for 30 minutes at room temperature. Eyes were washed in PBS for 30 minutes post-fixation. Tissue for cryosections was cryo-protected in 10% and then 20% sucrose in PBS for 1 hour each at 4 °C. The lenses were removed and the eye cups were placed in cryomold with Optimal Cutting Temperature media and frozen. Retinas were sectioned at 20µm using a cryostat. Slide mounted retinal sections were washed for 10 mins in PBS and blocked with 2% normal donkey serum, 0.2% TritonX-100 in PBS for 30 mins. Sections were incubated in primary antibody in blocking buffer overnight at 4 °C. Primary antibodies were used at the dilutions listed in Table 3. Sections were washed 3 times for 10 mins in PBS and incubated in secondary antibodies in blocking buffer for 2 hrs at room temperature. All secondary antibodies were used at a 1:500 dilution. Retinal sections were washed three times in PBS for 10 mins with DAPI (1:5000) included in the first wash step. Tissue was mounted in Fluoromount-G (Southern Biotech) and cover slipped for imaging.

Retina flat-mounts were fixed and washed as described above. The retinas were isolated from the eye cup and flattened by making 3-4 equally spaced incisions from the edge of the retina. Retina flat-mounts were post-fixed in 4% PFA in PBS for 10 minutes to help maintain their shape. Retinas were washed in blocking buffer (4% normal donkey serum, 0.2% TritonX-100) 3 times for 30 minutes. Retinas were incubated in primary antibody in blocking buffer for 3 days at room temperature. Following incubation of primary antibody, retinas were washed 3 times for 30 minutes in blocking buffer. Retinas were then incubated in secondary antibody overnight at room temperature. Next, retinas were washed in PBS three times for 30 minutes, flattened on slides, and cover slipped in Fluoromount-G for imaging.

### Image Acquisition

All retinal sections were imaged on a Zeiss Axio Imager M2 upright microscope equipped with an ApoTome2 using a 20X objective. Retinal flatmounts for mosaic analysis, single-cell morphology, and tracer coupling experiments were imaged on a Zeiss LSM 880 confocal microscope using a 40X objective. Images were acquired using the Zeiss Zen Imaging software for both microscopes.

### Co-localization analysis

All co-localization experiments used retinal sections from the *Gbx2*^*CreERT2-IRES-EGFP*^; *Rosa26*^*LSL-tdTomato*^ mouse line. Co-localization was determined by antibody labeling within the soma of the Gbx2+ amacrine cell bodies. Images were collected and analyzed as z-stacks to confirm the antibody labeling was within the soma of the correct z-depth. Offline analysis was completed using FIJI (Schindelin *et al.*, 2012).

### Mosaic cell spacing analysis

To analyze cell density and mosaic cell spacing, we used a 500µm x 500µm area of tissue from 4-8 locations within a single retina. Retina orientation was maintained to make spatial measurements in dorsal, ventral, temporal, and nasal areas of the retina. In addition, measurements in the peripheral and central regions of the retina made in ROIs ∼200µm from the peripheral edge or optic nerve head respectively. Cell counts and X-Y coordinates were measured offline in FIJI (Schindelin et al. 2012) and density recovery profiles were obtained by analysis completed in WinDRP (Rodieck, 1991).

### Neuron morphology analysis

For analysis of dendrite stratification in cross-section, we made measurements of fluorescent intensity along IPL depth using FIJI. These values were binned into 5% increments along the IPL depth using the FIJI plugin, IPLaminator (Li *et al.*, 2016). For analysis of dendritic arbor morphology in retinal flatmounts we analyzed isolated single cells labeled using either sparse expression in *Gbx2*^*CreERT2-IRES-EGFP*^; *Tigre*^*LSL-GFP*^ mice or by targeted cell fills using Alexa Fluor 488 hydrazide (Fisher Scientific). Offline tracing and analysis of dendritic arbors were made using the Filaments plugin in Imaris (Bitplane). Dendrite density (L/A) was calculated by dividing dendrite length (L) over dendrite area (A). Coverage factor was calculated by dividing the dendrite area over the cell density of Gbx2+ ACs in the INL and GCL for each subtype. Symmetry Index was calculated by subtracting the sums of missing dendrite coverage from 360 and then divided by 360, as previously described (Sun et al. 2013). Dendrite orientation was determined by the direction of the longest dendritic branch of a single cell and values were binned into 8 different groups based on the cardinal directions.

### Electrophysiology

Adult mice of either sex were dark adapted for 1-2 hours. Animals were anesthetized with isoflurane before being euthanized via cervical dislocation. Subsequent to enucleation, all procedures were performed under infrared illumination. The retina and attached pigment epithelium were dissected free from the sclera and placed in a recording chamber under a microscope and continuously perfused (5ml/min) with Ames medium maintained at 34°C. Cells were visualized though a 40x water-immersion objective and Dodt contrast illumination. Fluorescent Gbx2+ ACs were targeted under 2-photon guidance (excitation wavelength: 920 nm) with a Ti:sapphire laser (Chameleon ultra II; Coherent). Correct targeting was confirmed by visualizing the Alexa dye in intracellular solution fill the soma and processes of the target cell.

Patch electrodes were pulled from borosilicate glass to a final resistance of 8-12 MΩ. For voltage-clamp recordings, pipettes were filled with an intracellular solution containing the following (in mM): 125 Cs-methanesulphonate, 7 CsCl, 10 Na-HEPES, 3 phosphocreatine-Na_2_, 1 EGTA, 2 Mg-ATP, 1 Na-GTP, 0.1 Alexa Fluor 488 hydrazide, and 3 QX-314 chloride. The solution was adjusted to pH 7.35 using CsOH. Cesium was included in place of potassium to block voltage-gated potassium currents, thereby improving the voltage clamp at positive potentials. QX-314 was included to block voltage-gated sodium channels. For current-clamp recordings, all solution components were the same except potassium was used in place of cesium, and QX-314 was not included. Currents were sampled at 10 kHz and filtered at 2 kHz through the four-pole Bessel filter in an EPC-10 patch clamp amplifier (HEKA). Voltages were corrected for a liquid junction potential of -10mV.

Visual stimuli were produced using custom software based on PsychoPy routines (Peirce, 2007). The stimuli, generated on a Texas Instruments digital light projector (DLP; Lightcrafter 4500), were projected onto the photoreceptor layer through a 10x water immersion objective (0.3 NA, Olympus). The DLP intensity was linearized using a calibrated lookup table. DLP intensity was attenuated using neutral density filters to produce a grey adapting background flux of ∼3.4×10^5^ photons/µm^2^/s. Stimuli were first aligned to the receptive field center of each cell using a series of 100 x 1000 µm vertical and horizontal bars to locate the cell’s maximal response. All subsequent stimuli were centered on the coordinate of maximum response. Receptive field sizes were estimated from area-response data and fit to a difference of Gaussians function:

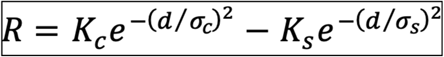

where *R* is the peak response evoked by a stimulus of diameter *d, K*_*c*_ and *K*_*s*_ are the amplitudes of the excitatory and inhibitory components, respectively, and *σ*_*c*_ and *σ*_*s*_ are their space constants.

### Pharmacological agents

Drugs were added to the perfusion solution. The following agents were used: L-(+)-2-amino-4-phophonobutyric acid (L-AP4; 20μM; Tocris Bioscience, catalog #0103), D-(-)-2-amino-5-phophonopentanoic acid (D-AP5; 50μM; Abcam Biochemicals, catalog #120003), 1-(4-aminophenyl)-3-methylcarbamyl-4-methyl-3,4-dihydro-7,8-meth-ylenedioxy-5H-2,3-benzodiazepine hydrochloride [GYKI-53655 (GYKI); 50μM;Tocris Bioscience catalog #2555), (*S*)-1-(2-Amino-2-carboxyethyl)-3-(2-carboxy-5-phenylthiophene-3-yl-methyl)-5-methylpyrimidine-2,4-dione (ACET; 1μM; Tocris Bioscience, catalog #2728).

### Data analysis

Light-evoked synaptic conductances were calculated as described previously (Taylor & Vaney, 2002). Briefly, current–voltage (I–V) relations were measured at 10 ms intervals over a range of voltage steps from -90 +50 mV in 20mV increments. The total light-evoked conductance was calculated as the difference between the I–V relation at each time point and the “leak” I–V relation measured just before the onset of the light stimulus. To avoid errors in calculating the net light-evoked currents due to a sloping baseline during positive voltage steps, a single exponential trend was subtracted from the current traces for each voltage step before the leak subtraction. The excitatory and inhibitory conductances could then be calculated at each time point using the observed I–V reversal potential along with the cation and chloride reversal potentials.

### Tracer coupling

*Gbx2*^*CreERT2-IRES-EGFP*^; *Rosa26*^*LSL-tdTomato*^ mice were used to target tdTomato positive neurons in the GCL for electroporation of Neurobiotin, as previously described (Kanjhan & Vaney, 2008; Sivyer & Vaney, 2010). In isolated preparations of retina bathed in Ames medium at room temperature, tdTomato positive neurons were lose seal patched with an intracellular solution containing (in mM): 120 K-gluconoate; 6 KCl;10 HEPES Na;10 Phosphocreatine-Na_2_; 60 Neurobiotin-Cl; 2 ATP; 0.5 GTP pH balanced to 7.2 with KOH. Once a seal was established the HEKA EPC-800 amplifier was switched to current clamp mode and ACs were electroporated by applying 0.5-1nA pulses for 0.5 s at 1 Hz. Following electroporation, retinae were incubated in Ames medium for 45 mins at 35 C before being transferred to room temperature Ames, mounted on nitrocellulose filter paper and fixed in cold paraformaldehyde in PBS for 30 mins. Retinas were then wash twice with 0.2% Triton in 1X PBS and incubated with 10 µg/mL Streptavidin Alexa Fluor 488 (Thermo Fisher) at room temperature overnight. Before imaging, stained retinas were washed three times in 1X PBS before being mounted on a glass coverslip.

### Retina Dissociation and FACS

Retinas from P6 or P7 mouse pups were dissected and isolated in 1X Hank’s Balanced Salt Solution (HBSS). Cells were dissociated into a single-cell suspension by incubating the retinas in 1mL of HBSS containing 10U of papain (Roche) and 200µM cysteine. Retinal tissue was incubated at 37 °C for 30 mins. After papain incubation, the tissue was pelleted using a bench top microfuge and washed twice with 1mL of HBSS. Cells were gently dissociated in 500µL of HBSS by flicking the tube. Dissociated cells were passed through a cell strainer (35µm nylon mesh) to remove any cell aggregates. DNase was added to the solution and the samples were placed on ice until sorted. Gbx2+ amacrine cells were sorted into two groups S3-cells that were both GFP+ and tdTomato+ and S5-cells that were only tdTomato+. All fluorescence activated cell sorting (FACS) was completed in the OHSU Flow Cytometry Core using a BD InFlux equipped with 488nm and 561nm laser lines. All cells were sorted directly into cell lysis buffer and RNA was isolated with Agilent Absolutely RNA Nanoprep Kit.

### RNAseq library preparation, sequencing, and analysis

The cDNA libraries for used for sequencing of 4 total RNA samples were synthesized using a SMART-Seq Ultra Low Input RNA kit (Takara) in the OHSU Massively Parallel Sequencing Shared Resource Core Facility. Two of the cDNA libraries contained tdTomato+/GFP+ cells (S3-Gbx2+ ACs) and two samples contained the cDNA libraries for tdTomato+ only cells (S5-Gbx2+ ACs). Quality and quantity of the cDNA libraries was determined on a Bioanalyzer. For multiplex sequencing, all four cDNA sample libraries were loaded on a single lane on a HiSeq 2500 sequencer (Illumina). Libraries were sequenced to a depth of 45-55 million reads per sample. Alignment rate of total reads was >97% across all samples.

Differential expression analysis was performed by the ONPRC Bioinformatics & Biostatistics Core. The quality of the raw sequencing files was evaluated using FastQC (Andrews, 2010) combined with MultiQC (http://multiqc.info/) (Ewels *et al.*, 2016). Trimmomatic was used to remove any remaining Illumina adapters (Bolger *et al.*, 2014). Reads were aligned to Ensembl’s GRCm38 along with its corresponding annotation, release 99. The program STAR (v2.7.3a) was used to align the reads to the genome (Dobin *et al.*, 2013). STAR has been shown to perform well compared to other RNA-seq aligners (Engstrom *et al.*, 2013). Since STAR utilizes the gene annotation file, it also calculated the number of reads aligned to each gene. RNA-SeQC (DeLuca *et al.*, 2012) and another round of MultiQC were utilized to ensure alignments were of sufficient quality.

Gene-level differential expression analysis was performed in open source software R (R Core Team, 2017). Gene-level raw counts were filtered to remove genes with extremely low counts in many samples following the published guidelines (Chen *et al.*, 2016), normalized using the trimmed mean of M-values method (TMM)(Robinson and Oshlack 2010), and transformed to log-counts per million with associated sample wise quality weight and observational precision weights using voom method (Law *et al.*, 2014). Gene-wise linear models comparing the cell types (tdTom+ vs. GFP+/tdTom+) were employed for differential expression analyses using limma with empirical Bayes moderation (Ritchie *et al.*, 2015) and false discovery rate (FDR) adjustment (Benjamini & Hochberg, 1995).

RNAseq datasets were organized and displayed into gene families with R Studio and the START app (Nelson *et al.*, 2017); and using the reference gene groups determined by the *HUGO Gene Nomenclature Committee at the European Bioinformatics Institute* (www.genenames.org). All graphs displaying RNAseq data were made using the Prism 8 Software (Graphpad Software, Inc.).

### Statistical analysis

For each experiment and time point a minimum of 3 retinas from three different mice were analyzed. For analysis of neuron morphology and tracer coupling, at least 5 neurons were analyzed from at least 3 animals. For all data sets, the variance was reported as mean ± SEM. Each data set was first tested for normality. Analysis between two groups was completed by using unpaired Student’s t-test (parametric) or Mann-Whitney U test (nonparametric). For analysis between more than two groups, we used either a one-way analysis of variance (ANOVA) with Tukey’s multiple comparison test (parametric) or Kruskal-Wallis with Dunn’s multiple comparison test (nonparametric). All statistical significance tests were completed using Prism 8 Software (Graphpad Software, Inc.) and Igor Pro 8.02 (WaveMetrics, Inc.) for electrophysiological analyses.

## ACKNOWLEDGEMENTS

We would like to thank the members of the Wright, Taylor, and Sivyer laboratories for their assistance and discussion throughout the course of this study. We also thank Dr. Kelly Monk and Dr. Martin Riccomagno for their comments on the manuscript. This work was supported by NIH grant R01 NS091027, a Whitehall Foundation Research grant, and an OHSU University Shared Resources Grant to K.M.W.; NIH grant F32 EY029974, the Collins Medical Trust, and the Knights Templar Eye Foundation to P.C.K.; NIH grants R01 EY022070 and P30 EY003176 to W.R.T.; NIH grant P30 EY010572 and unrestricted departmental funding from Research to Prevent Blindness (New York, NY) and support from the donors of National Glaucoma Research, a program of BrightFocus Foundation, to B.S. Confocal microscopy and analysis was performed in the OHSU Advance Light Microscopy Core supported by the NIH grant P30 NS061800. Cell sorting was performed in the OHSU Flow Cytometry Shared Resource and the core is supported by the Knight NCI Cancer Center Support Grant. Short read sequencing assays were performed by Dr. Amy Carlos and Dr. Robert Searles of the OHSU Massively Parallel Sequencing Shared Resource. The authors acknowledge the support of Dr. Suzanne Fei and Dr. Lina Gao of the Oregon National Primate Research Center Bioinformatics & Biostatistics Core, which is funded in part by NIH grant OD P51 OD011092.

---

**Figure 1 – Supplement 1.**
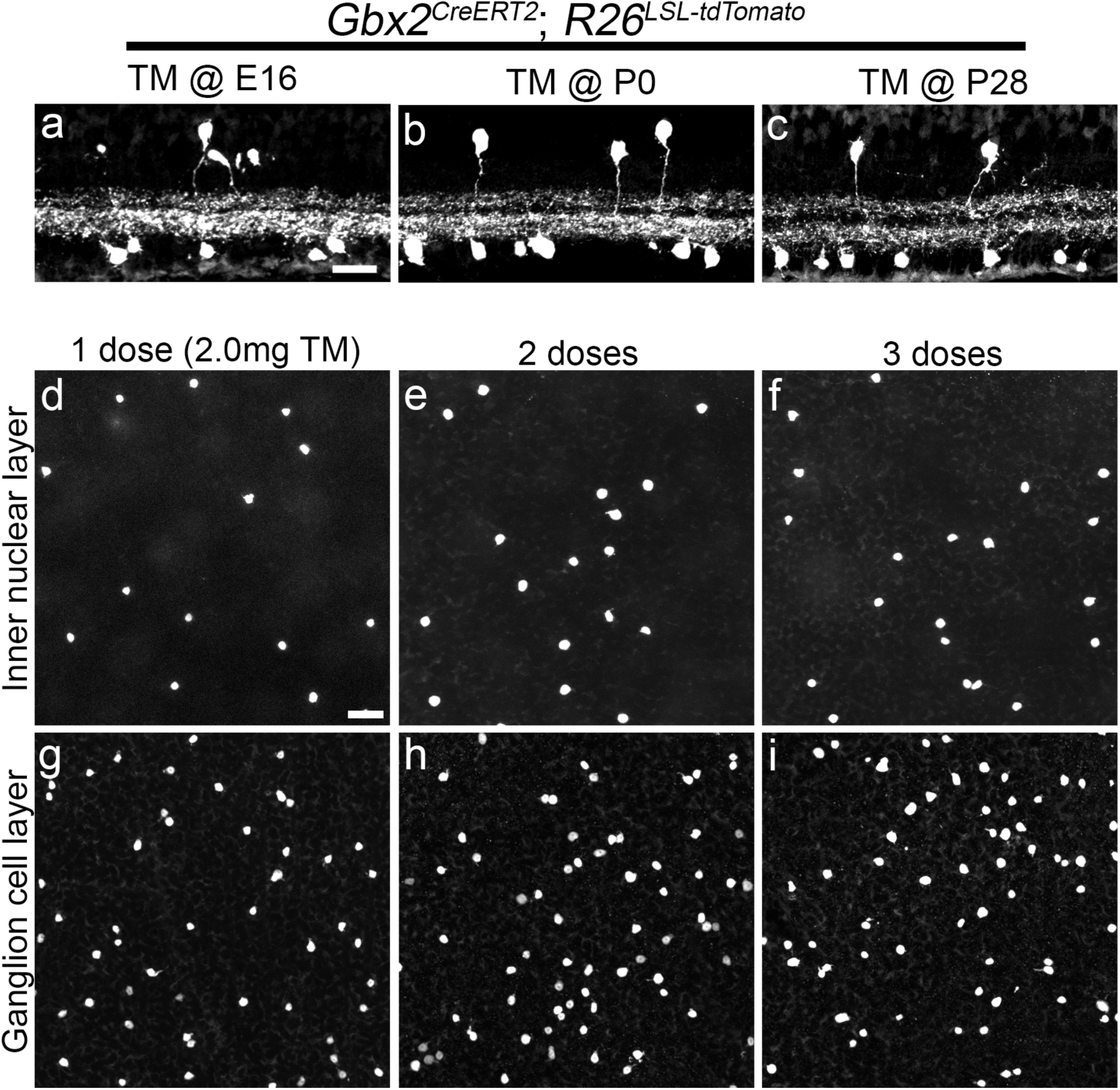
*Gbx2*^*CreERT2-IRES-EGFP*^ expression is consistent throughout the mouse development and adulthood. **(a-c)** Adult (P35) Retinal cross-sections from *Gbx2*^*CreERT2-IRES-EGFP*^; *R26*^*LSL-tdTom*^ mice dosed with of Tamoxifen (60mg/kg) at **(a)** E16, **(b)** P0, **(c)** P28. (**d-i**) Z-projections through the cell bodies of the inner nuclear layer (**d-f**) and ganglion cell layer (**g-i**) from retinal flatmounts of *Gbx2*^*CreERT2-IRES-EGFP*^; *R26*^*LSL-tdTom*^ mice administered 2.0mg/day tamoxifen for (**d**,**g**) 1 day (dose), (**e**,**h**) 2 days, (**f-i**) 3 days. Scale bar, 25μm in (**a**) and (**d**).

**Figure 1 – Supplement 2.**
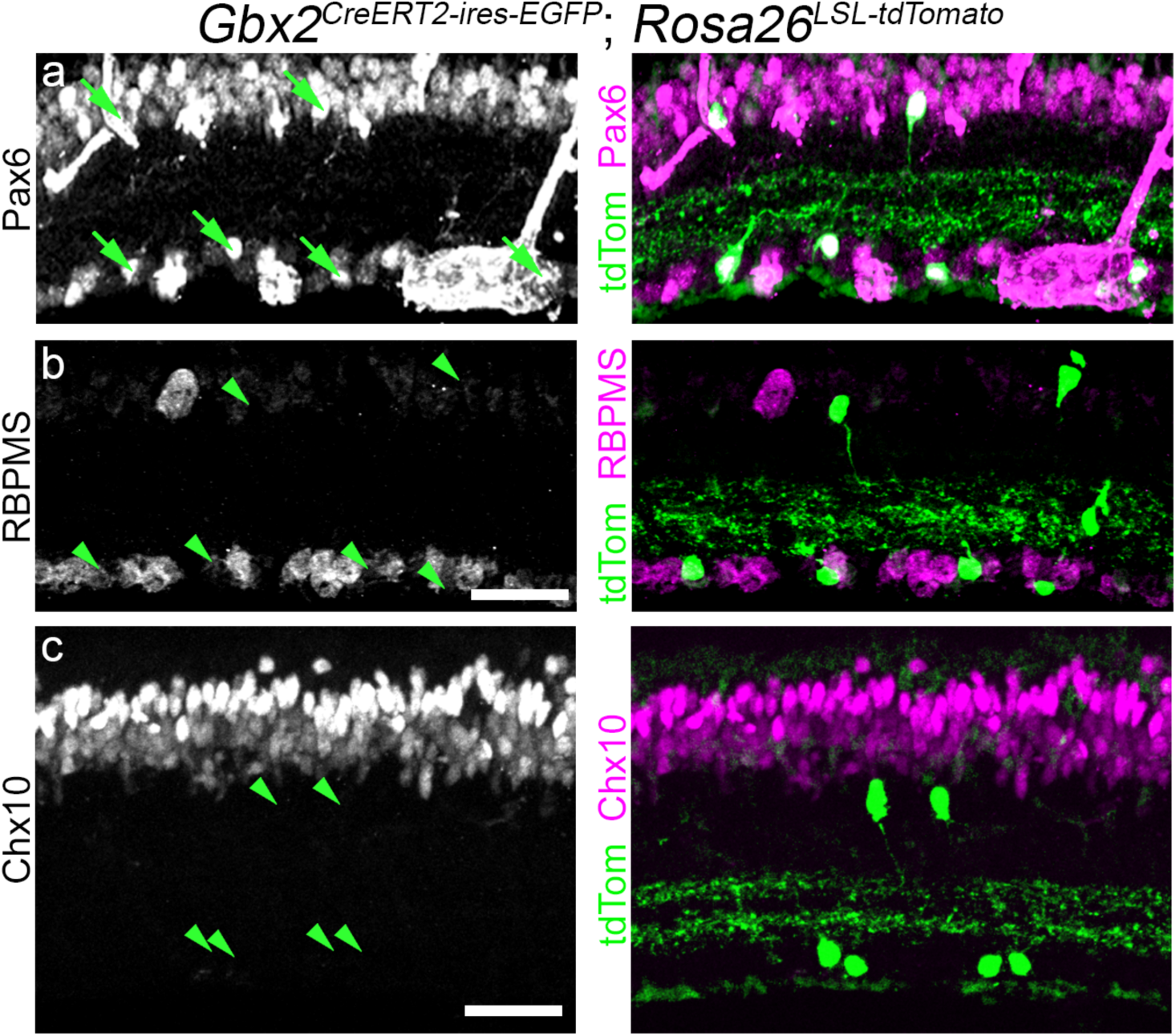
Gbx2+ retinal neurons are amacrine cells. Cross-sections of an adult retina from a *Gbx2*^*CreERT2-IRES-EGFP*^; *Rosa26*^*LSL-tdTom*^ mouse labeling the total Gbx2+ AC population. Left: Neurotransmitter markers, (**a**) Pax6, (**b**) RBPMS, and (**c**) Chx10, label amacrine cells, retinal ganglion cells, and bipolar cells in the inner nuclear layer and ganglion cell layer, respectively. Right: Merged images showing both the select neurotransmitter marker (magenta) and Gbx2+ ACs (green). Arrows denote colocalization between the cell marker and Gbx2+ ACs, and arrowheads denote Gbx2+ ACs that do not colocalize with the specific cell marker. Scale bar, 25 µm.

**Figure 1 – Supplement 3.**
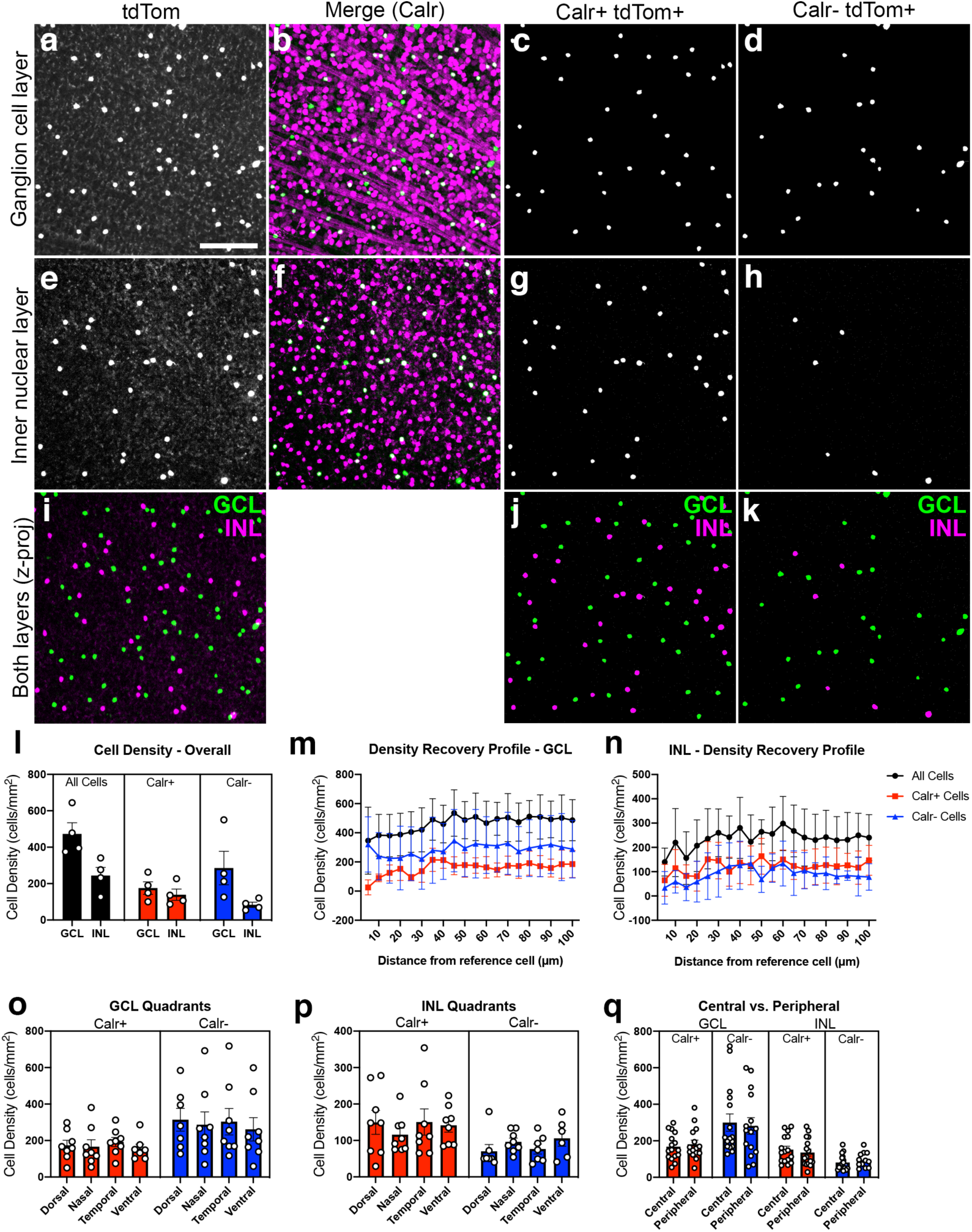
Gbx2+ AC subpopulations have consistent density and spacing across the retina. **(a)**Tdtomato expressing Gbx2+ ACs in the ganglion cell layer (GCL) in a retina flatmount from a *Gbx2*^*CreER*^; *R26*^*LSL-tdTomato*^ mouse. **(b)** Gbx2+ ACs (green) immunolabeled for calretinin (Calr, magenta) in the GCL. **(c-d)** Masked image of cell bodies of Gbx2+ neurons **(c)** Calretinin+ and **(d)** Calretinin-from the image in **(b). (e)** TdTomato expressing Gbx2+ ACs in the inner nuclear layer (INL). **(f)** Gbx2+ ACs (green) immunolabeled for calretinin (magenta) in the INL. **(g-h)** Masked image of cell bodies of Gbx2+ neurons **(c)** Calr+ and **(d)** Calr-from the image in **(f). (i-k)** Z-projection through the GCL and INL (pseudocolored green and magenta, respectively) for **(i)** all Gbx2+ ACs, **(j)** Calr+ Gbx2+ ACs, and **(k)** Calr-Gbx2+ ACs. **(l)** Quantification of the cell density of Calr+ and Calr-Gbx2+ ACs in the GCL and INL (n= 24 measurements from 4 mice). **(m-n)** The density recovery profile (DRP) of Calr+ and Calr-Gbx2 ACs in the **(m)** GCL and **(n)** INL (n= 32 measurements from 4 mice, respectively). **(o-q)** The cell densities in the four quadrants of the retina of Gbx2+ AC in the **(o)** GCL and **(p)** INL (n= 32 measurements from 4 mice, respectively). **(q)** The cell density of Gbx2+ AC populations in the central and peripheral retina (n= 32 measurements from 4 mice, respectively). Scale bar, 50 μm in **(a)**.

**Figure 2 – Supplement 1.**
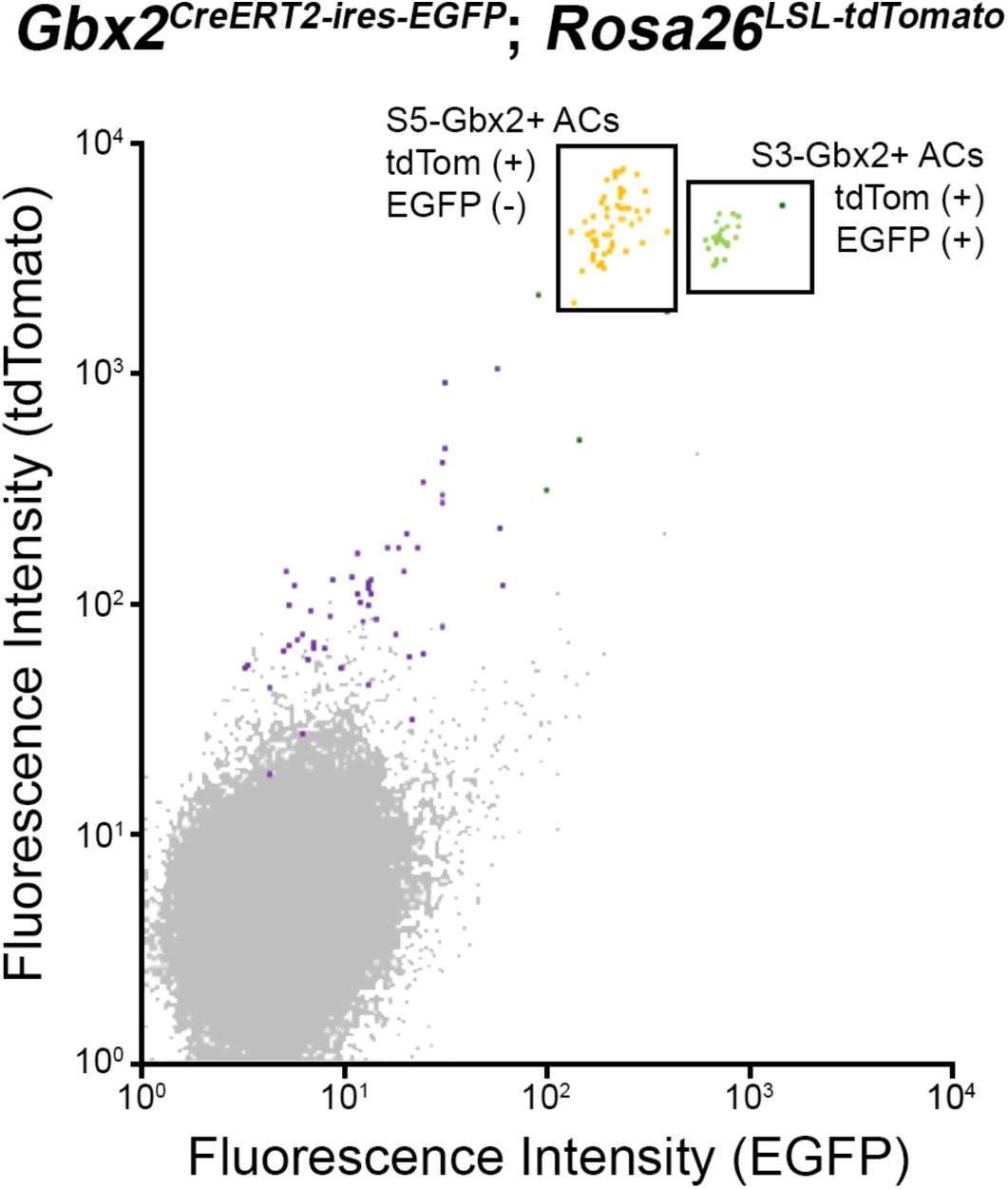
Flow cytometry plot of dissociated retinal neurons isolated from a *Gbx2*^*CreERT2-IRES-EGFP*^; *Rosa26*^*LSL-tdTomato*^ mouse. Using fluorescence-activated cell sorting, Gbx2+ ACs (tdTomato+) from P8 retina were isolated and separated into the S5-Gbx2+ ACs (tdTomato+, EGFP^low^) and the S3-Gbx2+ ACs (tdTomato+, EGFP^high^) groups for bulk RNA sequencing.

**Figure 3 – Supplement 1.**
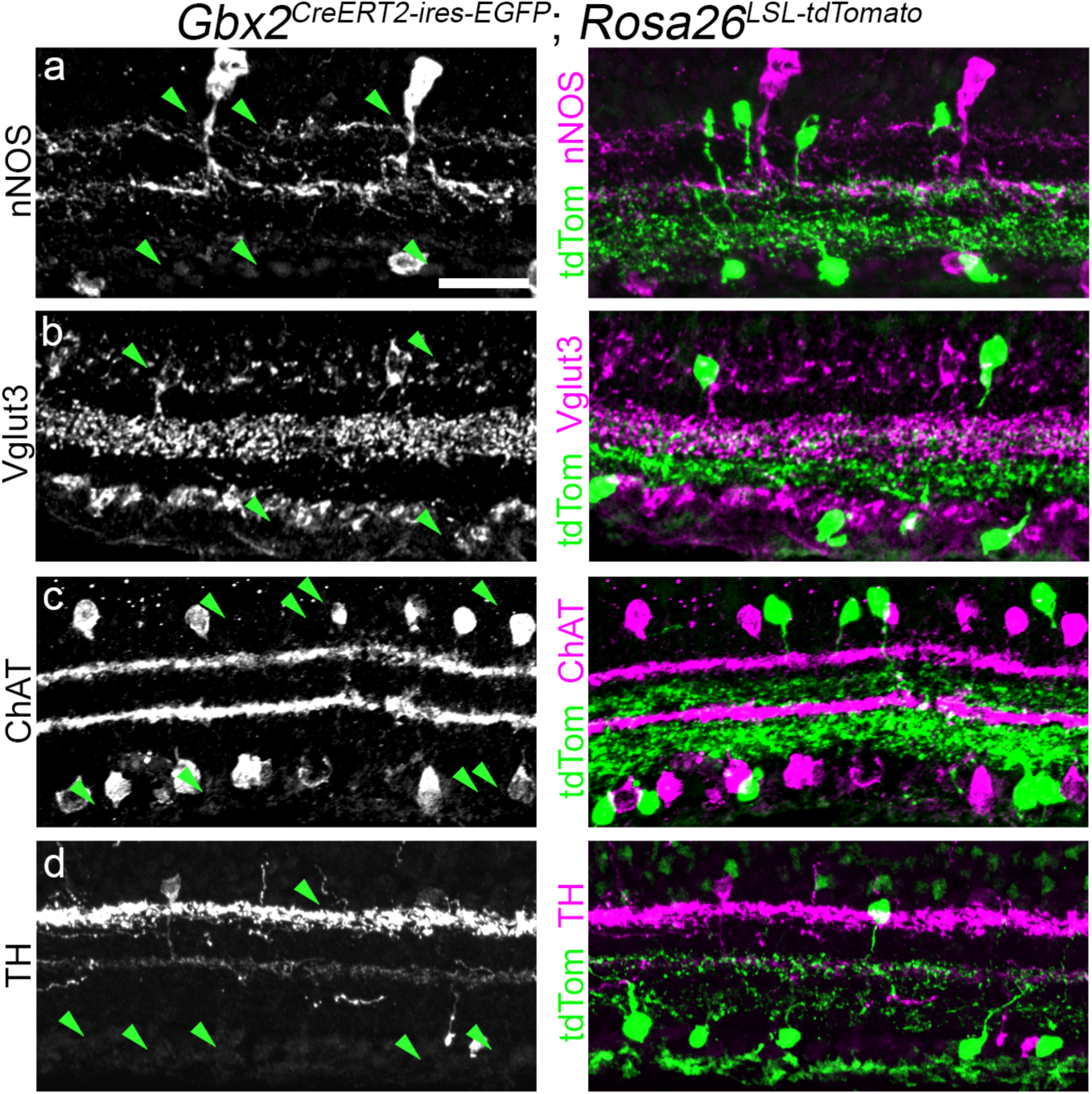
Gbx2+ ACs do not colocalize with many canonical neurotransmitter cell markers. Cross-sections of an adult retina from a *Gbx2*^*CreERT2-IRES-EGFP*^; *Rosa26*^*LSL-tdTom*^ mouse labeling the total Gbx2+ AC population (high-TM, 2.0mg tamoxifen). Left: Left, retinal sections from a *Gbx2*^*CreERT2-IRES-EGFP*^; *Rosa26*^*LSL-tdTom*^ mouse immunolabeled with (**a**) neuronal nitric oxide synthase (nNOS), (**b**) vesicular glutamate transporter 3 (Vglut3), (**c**) choline acetyl transferase transporter (ChAT), and (**d**) tyrosine hydroxylase (TH). Right: Merged images of Gbx2+ ACs (green) and the neurotransmitter marker (magenta). Arrows denote colocalization between the cell marker and Gbx2+ ACs, and arrowheads denote Gbx2+ ACs that do not colocalize with the specific cell marker. Scale bar, 25 µm.

**Figure 4 – Supplement 1.**
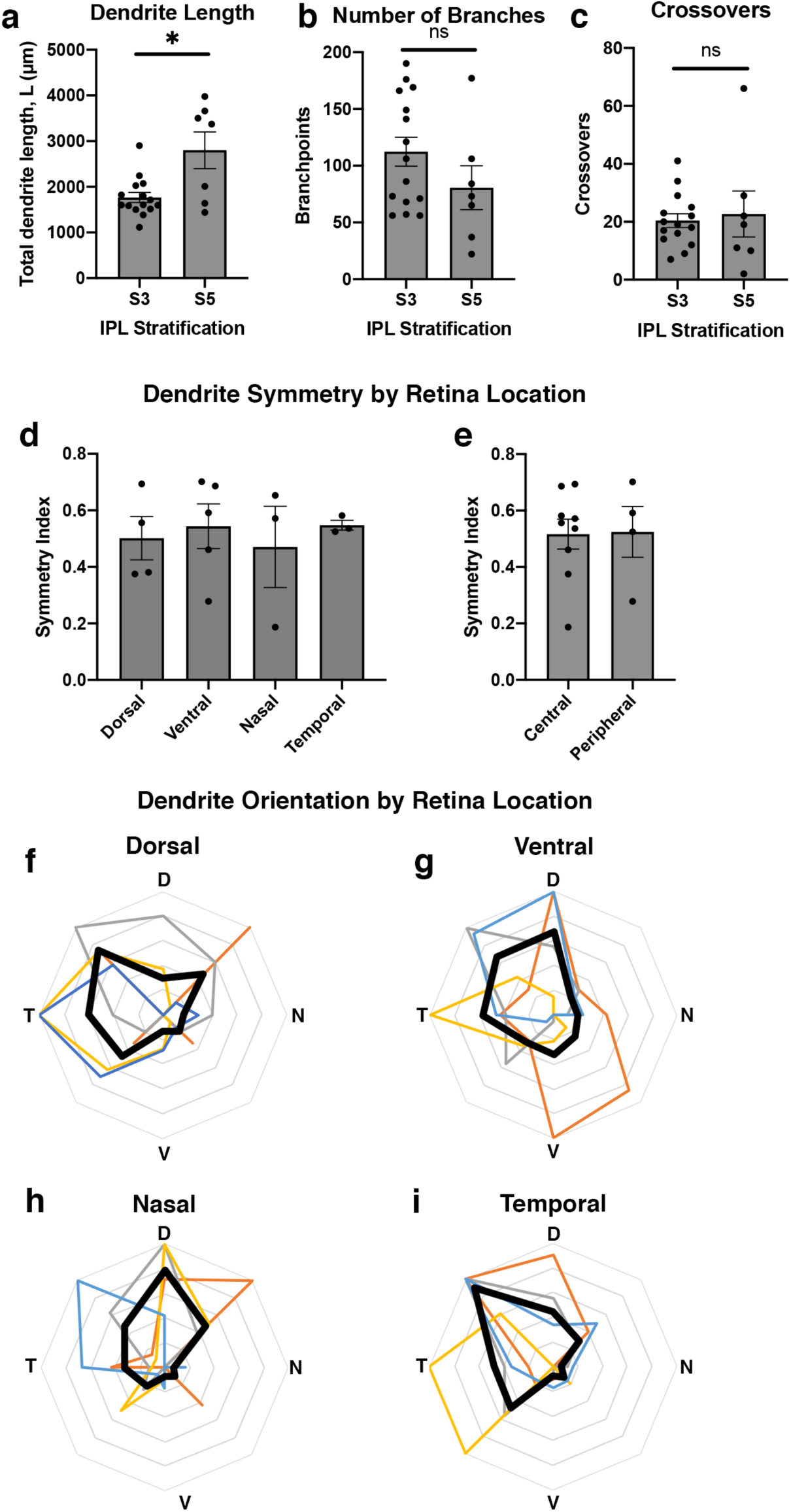
Dendritic morphology and orientation of Gbx2+ ACs by retina location. **(a-c)** S3- and S5-Gbx2+ AC morphology in (**a**) total dendrite length, **(b)** number of branches, **(c)** dendrite branch self-crossover. n=15, 7 cells for S3 and S5 respectively. **(d, e)** S3-stratifying Gbx2+ ACs show a similar dendrite asymmetry in **(d)** each retinal quadrant (n=3-5 retinas per quadrant) and **(e)** between central and peripheral retina (n=4, 9 retinas respectively). **(f-i)** A polar plot of dendrite orientation of S3-targeting Gbx2+ ACs in each retinal quadrant; black trace represents the mean and colored traces represent neurons quantified from a single retina (n >20 neurons per retina, n= 4 retinas). D, dorsal; V, ventral; N, nasal; T, temporal. *p<0.05 by an unpaired t-test with a Welch’s correction.

## BIBLIOGRAPHY

Andrews, S. 2010 FastQC: A quality control tool for high throughput sequence data. <http://www.bioinformatics.babraham.ac.uk/projects/fastqc/>

Badea, T.C. & Nathans, J. (2004) Quantitative analysis of neuronal morphologies in the mouse retina visualized by using a genetically directed reporter. J Comp Neurol, 480, 331–351.

Baden, T., Berens, P., Franke, K., Roman Roson, M., Bethge, M. & Euler, T. (2016) The functional diversity of retinal ganglion cells in the mouse. Nature, 529, 345-350. PMC4724341

Benjamini, Y. & Hochberg, Y. (1995) Controlling the False Discovery Rate: A Practical and Powerful Approach to Multiple Testing. Journal of the Royal Statistical Society. Series B (Methodological), 57, 289–300.

Bloomfield, S.A. & Volgyi, B. (2009) The diverse functional roles and regulation of neuronal gap junctions in the retina. Nat Rev Neurosci, 10, 495-506. PMC3381350

Bolger, A.M., Lohse, M. & Usadel, B. (2014) Trimmomatic: a flexible trimmer for Illumina sequence data. Bioinformatics, 30, 2114-2120. PMC4103590

Brecha, N., Johnson, D., Peichl, L. & Wassle, H. (1988) Cholinergic amacrine cells of the rabbit retina contain glutamate decarboxylase and gamma-aminobutyrate immunoreactivity. Proc Natl Acad Sci U S A, 85, 6187-6191. PMC281930

Chatterjee, M., Li, K., Chen, L., Maisano, X., Guo, Q., Gan, L. & Li, J.Y. (2012) Gbx2 regulates thalamocortical axon guidance by modifying the LIM and Robo codes. Development, 139, 4633-4643. PMC3509725

Chen, L., Chatterjee, M. & Li, J.Y. (2010) The mouse homeobox gene Gbx2 is required for the development of cholinergic interneurons in the striatum. J Neurosci, 30, 14824-14834. PMC3071646

Chen, L., Guo, Q. & Li, J.Y. (2009) Transcription factor Gbx2 acts cell-nonautonomously to regulate the formation of lineage-restriction boundaries of the thalamus. Development, 136, 1317–1326. PMC2687463

Chen, Y., Lun, A.T. & Smyth, G.K. (2016) From reads to genes to pathways: differential expression analysis of RNA-Seq experiments using Rsubread and the edgeR quasi-likelihood pipeline. F1000Res, 5, 1438. PMC4934518

Cherry, T.J., Trimarchi, J.M., Stadler, M.B. & Cepko, C.L. (2009) Development and diversification of retinal amacrine interneurons at single cell resolution. Proc Natl Acad Sci U S A, 106, 9495-9500. PMC2686638

Daigle, T.L., Madisen, L., Hage, T.A., Valley, M.T., Knoblich, U., Larsen, R.S., Takeno, M.M., Huang, L., Gu, H., Larsen, R., Mills, M., Bosma-Moody, A., Siverts, L.A., Walker, M., Graybuck, L.T., Yao, Z., Fong, O., Nguyen, T.N., Garren, E., Lenz, G.H., Chavarha, M., Pendergraft, J., Harrington, J., Hirokawa, K.E., Harris, J.A., Nicovich, P.R., McGraw, M.J., Ollerenshaw, D.R., Smith, K.A., Baker, C.A., Ting, J.T., Sunkin, S.M., Lecoq, J., Lin, M.Z., Boyden, E.S., Murphy, G.J., da Costa, N.M., Waters, J., Li, L., Tasic, B. & Zeng, H. (2018) A Suite of Transgenic Driver and Reporter Mouse Lines with Enhanced Brain-Cell-Type Targeting and Functionality. Cell, 174, 465-480 e422. PMC6086366

DeLuca, D.S., Levin, J.Z., Sivachenko, A., Fennell, T., Nazaire, M.D., Williams, C., Reich, M., Winckler, W. & Getz, G. (2012) RNA-SeQC: RNA-seq metrics for quality control and process optimization. Bioinformatics, 28, 1530-1532. PMC3356847

Diamond, J.S. (2017) Inhibitory Interneurons in the Retina: Types, Circuitry, and Function. Annu Rev Vis Sci, 3, 1–24.

Dobin, A., Davis, C.A., Schlesinger, F., Drenkow, J., Zaleski, C., Jha, S., Batut, P., Chaisson, M. & Gingeras, T.R. (2013) STAR: ultrafast universal RNA-seq aligner. Bioinformatics, 29, 15-21. PMC3530905

Engstrom, P.G., Steijger, T., Sipos, B., Grant, G.R., Kahles, A., Ratsch, G., Goldman, N., Hubbard, T.J., Harrow, J., Guigo, R., Bertone, P. & Consortium, R. (2013) Systematic evaluation of spliced alignment programs for RNA-seq data. Nat Methods, 10, 1185-1191. PMC4018468

Estevez, M.E., Fogerson, P.M., Ilardi, M.C., Borghuis, B.G., Chan, E., Weng, S., Auferkorte, O.N., Demb, J.B. & Berson, D.M. (2012) Form and function of the M4 cell, an intrinsically photosensitive retinal ganglion cell type contributing to geniculocortical vision. J Neurosci, 32, 13608-13620. PMC3474539

Ewels, P., Magnusson, M., Lundin, S. & Kaller, M. (2016) MultiQC: summarize analysis results for multiple tools and samples in a single report. Bioinformatics, 32, 3047-3048. PMC5039924

Hampson, E.C., Vaney, D.I. & Weiler, R. (1992) Dopaminergic modulation of gap junction permeability between amacrine cells in mammalian retina. J Neurosci, 12, 4911-4922. PMC6575774

Haverkamp, S. & Wassle, H. (2004) Characterization of an amacrine cell type of the mammalian retina immunoreactive for vesicular glutamate transporter 3. J Comp Neurol, 468, 251–263.

He, L., Gulyanon, S., Mihovilovic Skanata, M., Karagyozov, D., Heckscher, E.S., Krieg, M., Tsechpenakis, G., Gershow, M. & Tracey, W.D., Jr. (2019) Direction Selectivity in Drosophila Proprioceptors Requires the Mechanosensory Channel Tmc. Curr Biol, 29, 945-956 e943. PMC6884367

Helmstaedter, M., Briggman, K.L., Turaga, S.C., Jain, V., Seung, H.S. & Denk, W. (2013) Connectomic reconstruction of the inner plexiform layer in the mouse retina. Nature, 500, 168–174.

Jacoby, J. & Schwartz, G.W. (2018) Typology and Circuitry of Suppressed-by-Contrast Retinal Ganglion Cells. Front Cell Neurosci, 12, 269. PMC6119723

Jacoby, J., Zhu, Y., DeVries, S.H. & Schwartz, G.W. (2015) An Amacrine Cell Circuit for Signaling Steady Illumination in the Retina. Cell Rep, 13, 2663-2670. PMC4698005

Johnson, J., Sherry, D.M., Liu, X., Fremeau, R.T., Jr., Seal, R.P., Edwards, R.H. & Copenhagen, D.R. (2004) Vesicular glutamate transporter 3 expression identifies glutamatergic amacrine cells in the rodent retina. J Comp Neurol, 477, 386-398. PMC2586940

Kanjhan, R. & Vaney, D.I. (2008) Semi-loose seal Neurobiotin electroporation for combined structural and functional analysis of neurons. Pflugers Arch, 457, 561–568.

Kay, J.N., Voinescu, P.E., Chu, M.W. & Sanes, J.R. (2011) Neurod6 expression defines new retinal amacrine cell subtypes and regulates their fate. Nat Neurosci, 14, 965-972. PMC3144989

Keeley, P.W., Eglen, S.J. & Reese, B.E. (2020) From random to regular: Variation in the patterning of retinal mosaics. J Comp Neurol.

Kim, I.J., Zhang, Y., Yamagata, M., Meister, M. & Sanes, J.R. (2008) Molecular identification of a retinal cell type that responds to upward motion. Nature, 452, 478–482.

Kim, J.I., Ganesan, S., Luo, S.X., Wu, Y.W., Park, E., Huang, E.J., Chen, L. & Ding, J.B. (2015) Aldehyde dehydrogenase 1a1 mediates a GABA synthesis pathway in midbrain dopaminergic neurons. Science, 350, 102-106. PMC4725325

Krishnaswamy, A., Yamagata, M., Duan, X., Hong, Y.K. & Sanes, J.R. (2015) Sidekick 2 directs formation of a retinal circuit that detects differential motion. Nature, 524, 466-470. PMC4552609

Law, C.W., Chen, Y., Shi, W. & Smyth, G.K. (2014) voom: Precision weights unlock linear model analysis tools for RNA-seq read counts. Genome Biol, 15, R29. PMC4053721

Lee, S., Chen, L., Chen, M., Ye, M., Seal, R.P. & Zhou, Z.J. (2014) An unconventional glutamatergic circuit in the retina formed by vGluT3 amacrine cells. Neuron, 84, 708-715. PMC4254642

Lefebvre, J.L., Sanes, J.R. & Kay, J.N. (2015) Development of dendritic form and function. Annu Rev Cell Dev Biol, 31, 741–777.

Li, S., Woodfin, M., Long, S.S. & Fuerst, P.G. (2016) IPLaminator: an ImageJ plugin for automated binning and quantification of retinal lamination. BMC Bioinformatics, 17, 36. PMC4715356

Lin, B. & Masland, R.H. (2006) Populations of wide-field amacrine cells in the mouse retina. J Comp Neurol, 499, 797–809.

Liu, J., Reggiani, J.D.S., Laboulaye, M.A., Pandey, S., Chen, B., Rubenstein, J.L.R., Krishnaswamy, A. & Sanes, J.R. (2018) Tbr1 instructs laminar patterning of retinal ganglion cell dendrites. Nat Neurosci, 21, 659-670. PMC5920715

Liu, J. & Sanes, J.R. (2017) Cellular and Molecular Analysis of Dendritic Morphogenesis in a Retinal Cell Type That Senses Color Contrast and Ventral Motion. J Neurosci, 37, 12247-12262. PMC5729193

Luu, B., Ellisor, D. & Zervas, M. (2011) The lineage contribution and role of Gbx2 in spinal cord development. PLoS One, 6, e20940. PMC3116860

MacNeil, M.A., Heussy, J.K., Dacheux, R.F., Raviola, E. & Masland, R.H. (1999) The shapes and numbers of amacrine cells: matching of photofilled with Golgi-stained cells in the rabbit retina and comparison with other mammalian species. J Comp Neurol, 413, 305–326.

MacNeil, M.A. & Masland, R.H. (1998) Extreme diversity among amacrine cells: implications for function. Neuron, 20, 971–982.

Macosko, E.Z., Basu, A., Satija, R., Nemesh, J., Shekhar, K., Goldman, M., Tirosh, I., Bialas, A.R., Kamitaki, N., Martersteck, E.M., Trombetta, J.J., Weitz, D.A., Sanes, J.R., Shalek, A.K., Regev, A. & McCarroll, S.A. (2015) Highly Parallel Genome-wide Expression Profiling of Individual Cells Using Nanoliter Droplets. Cell, 161, 1202-1214. PMC4481139

Madisen, L., Zwingman, T.A., Sunkin, S.M., Oh, S.W., Zariwala, H.A., Gu, H., Ng, L.L., Palmiter, R.D., Hawrylycz, M.J., Jones, A.R., Lein, E.S. & Zeng, H. (2010) A robust and high-throughput Cre reporting and characterization system for the whole mouse brain. Nat Neurosci, 13, 133-140. PMC2840225

Mallika, C., Guo, Q. & Li, J.Y. (2015) Gbx2 is essential for maintaining thalamic neuron identity and repressing habenular characters in the developing thalamus. Dev Biol, 407, 26-39. PMC4641819

Nelson, J.W., Sklenar, J., Barnes, A.P. & Minnier, J. (2017) The START App: a web-based RNAseq analysis and visualization resource. Bioinformatics, 33, 447-449. PMC6075080

Newkirk, G.S., Hoon, M., Wong, R.O. & Detwiler, P.B. (2013) Inhibitory inputs tune the light response properties of dopaminergic amacrine cells in mouse retina. J Neurophysiol, 110, 536-552. PMC3727066

O’Brien, J. & Bloomfield, S.A. (2018) Plasticity of Retinal Gap Junctions: Roles in Synaptic Physiology and Disease. Annu Rev Vis Sci, 4, 79–100.

Park, S.J.H., Pottackal, J., Ke, J.B., Jun, N.Y., Rahmani, P., Kim, I.J., Singer, J.H. & Demb, J.B. (2018) Convergence and Divergence of CRH Amacrine Cells in Mouse Retinal Circuitry. J Neurosci, 38, 3753-3766. PMC5895998

Peirce, J.W. (2007) PsychoPy--Psychophysics software in Python. J Neurosci Methods, 162, 8-13. PMC2018741

Peng, Y.R., James, R.E., Yan, W., Kay, J.N., Kolodkin, A.L. & Sanes, J.R. (2020) Binary Fate Choice between Closely Related Interneuronal Types Is Determined by a Fezf1-Dependent Postmitotic Transcriptional Switch. Neuron, 105, 464-474 e466. PMC7007373

Peng, Y.R., Shekhar, K., Yan, W., Herrmann, D., Sappington, A., Bryman, G.S., van Zyl, T., Do, M.T.H., Regev, A. & Sanes, J.R. (2019) Molecular Classification and Comparative Taxonomics of Foveal and Peripheral Cells in Primate Retina. Cell, 176, 1222-1237 e1222. PMC6424338

Peng, Y.R., Tran, N.M., Krishnaswamy, A., Kostadinov, D., Martersteck, E.M. & Sanes, J.R. (2017) Satb1 Regulates Contactin 5 to Pattern Dendrites of a Mammalian Retinal Ganglion Cell. Neuron, 95, 869-883 e866. PMC5575751

Pitulescu, M.E., Schmidt, I., Benedito, R. & Adams, R.H. (2010) Inducible gene targeting in the neonatal vasculature and analysis of retinal angiogenesis in mice. Nat Protoc, 5, 1518–1534.

Rheaume, B.A., Jereen, A., Bolisetty, M., Sajid, M.S., Yang, Y., Renna, K., Sun, L., Robson, P. & Trakhtenberg, E.F. (2018) Single cell transcriptome profiling of retinal ganglion cells identifies cellular subtypes. Nat Commun, 9, 2759. PMC6050223

Ritchie, M.E., Phipson, B., Wu, D., Hu, Y., Law, C.W., Shi, W. & Smyth, G.K. (2015) limma powers differential expression analyses for RNA-sequencing and microarray studies. Nucleic Acids Res, 43, e47. PMC4402510

Rodieck, R.W. (1991) The density recovery profile: a method for the analysis of points in the plane applicable to retinal studies. Vis Neurosci, 6, 95–111.

Roska, B. & Werblin, F. (2001) Vertical interactions across ten parallel, stacked representations in the mammalian retina. Nature, 410, 583–587.

Sanes, J.R. & Masland, R.H. (2015) The types of retinal ganglion cells: current status and implications for neuronal classification. Annu Rev Neurosci, 38, 221–246.

Saunders, A., Macosko, E.Z., Wysoker, A., Goldman, M., Krienen, F.M., de Rivera, H., Bien, E., Baum, M., Bortolin, L., Wang, S., Goeva, A., Nemesh, J., Kamitaki, N., Brumbaugh, S., Kulp, D. & McCarroll, S.A. (2018) Molecular Diversity and Specializations among the Cells of the Adult Mouse Brain. Cell, 174, 1015-1030 e1016. PMC6447408

Schindelin, J., Arganda-Carreras, I., Frise, E., Kaynig, V., Longair, M., Pietzsch, T., Preibisch, S., Rueden, C., Saalfeld, S., Schmid, B., Tinevez, J.Y., White, D.J., Hartenstein, V., Eliceiri, K., Tomancak, P. & Cardona, A. (2012) Fiji: an open-source platform for biological-image analysis. Nat Methods, 9, 676-682. PMC3855844

Schmidt, T.M., Chen, S.K. & Hattar, S. (2011) Intrinsically photosensitive retinal ganglion cells: many subtypes, diverse functions. Trends Neurosci, 34, 572-580. PMC3200463

Schwab, M.H., Bartholomae, A., Heimrich, B., Feldmeyer, D., Druffel-Augustin, S., Goebbels, S., Naya, F.J., Zhao, S., Frotscher, M., Tsai, M.J. & Nave, K.A. (2000) Neuronal basic helix-loop-helix proteins (NEX and BETA2/Neuro D) regulate terminal granule cell differentiation in the hippocampus. J Neurosci, 20, 3714-3724. PMC6772686

Shekhar, K., Lapan, S.W., Whitney, I.E., Tran, N.M., Macosko, E.Z., Kowalczyk, M., Adiconis, X., Levin, J.Z., Nemesh, J., Goldman, M., McCarroll, S.A., Cepko, C.L., Regev, A. & Sanes, J.R. (2016) Comprehensive Classification of Retinal Bipolar Neurons by Single-Cell Transcriptomics. Cell, 166, 1308-1323 e1330. PMC5003425

Siegert, S., Scherf, B.G., Del Punta, K., Didkovsky, N., Heintz, N. & Roska, B. (2009) Genetic address book for retinal cell types. Nat Neurosci, 12, 1197–1204.

Sivyer, B. & Vaney, D.I. (2010) Dendritic morphology and tracer-coupling pattern of physiologically identified transient uniformity detector ganglion cells in rabbit retina. Vis Neurosci, 27, 159–170.

Sonoda, T., Okabe, Y. & Schmidt, T.M. (2020) Overlapping morphological and functional properties between M4 and M5 intrinsically photosensitive retinal ganglion cells. J Comp Neurol, 528, 1028-1040. PMC7007370

Tasic, B., Yao, Z., Graybuck, L.T., Smith, K.A., Nguyen, T.N., Bertagnolli, D., Goldy, J., Garren, E., Economo, M.N., Viswanathan, S., Penn, O., Bakken, T., Menon, V., Miller, J., Fong, O., Hirokawa, K.E., Lathia, K., Rimorin, C., Tieu, M., Larsen, R., Casper, T., Barkan, E., Kroll, M., Parry, S., Shapovalova, N.V., Hirschstein, D., Pendergraft, J., Sullivan, H.A., Kim, T.K., Szafer, A., Dee, N., Groblewski, P., Wickersham, I., Cetin, A., Harris, J.A., Levi, B.P., Sunkin, S.M., Madisen, L., Daigle, T.L., Looger, L., Bernard, A., Phillips, J., Lein, E., Hawrylycz, M., Svoboda, K., Jones, A.R., Koch, C. & Zeng, H. (2018) Shared and distinct transcriptomic cell types across neocortical areas. Nature, 563, 72-78. PMC6456269

Taylor, W.R. & Vaney, D.I. (2002) Diverse synaptic mechanisms generate direction selectivity in the rabbit retina. J Neurosci, 22, 7712-7720. PMC6757986

Tran, N.M., Shekhar, K., Whitney, I.E., Jacobi, A., Benhar, I., Hong, G., Yan, W., Adiconis, X., Arnold, M.E., Lee, J.M., Levin, J.Z., Lin, D., Wang, C., Lieber, C.M., Regev, A., He, Z. & Sanes, J.R. (2019) Single-Cell Profiles of Retinal Ganglion Cells Differing in Resilience to Injury Reveal Neuroprotective Genes. Neuron, 104, 1039-1055 e1012. PMC6923571

Tritsch, N.X., Ding, J.B. & Sabatini, B.L. (2012) Dopaminergic neurons inhibit striatal output through non-canonical release of GABA. Nature, 490, 262-266. PMC3944587

Tritsch, N.X., Oh, W.J., Gu, C. & Sabatini, B.L. (2014) Midbrain dopamine neurons sustain inhibitory transmission using plasma membrane uptake of GABA, not synthesis. Elife, 3, e01936. PMC4001323

Vaney, D.I. (1991) Many diverse types of retinal neurons show tracer coupling when injected with biocytin or Neurobiotin. Neurosci Lett, 125, 187–190.

Vaney, D.I. & Weiler, R. (2000) Gap junctions in the eye: evidence for heteromeric, heterotypic and mixed-homotypic interactions. Brain Res Brain Res Rev, 32, 115–120.

Vaney, D.I. & Young, H.M. (1988) GABA-like immunoreactivity in cholinergic amacrine cells of the rabbit retina. Brain Res, 438, 369–373.

Whitney, I.E., Keeley, P.W., St John, A.J., Kautzman, A.G., Kay, J.N. & Reese, B.E. (2014) Sox2 regulates cholinergic amacrine cell positioning and dendritic stratification in the retina. J Neurosci, 34, 10109-10121. PMC4107400

Yadav, S.C., Tetenborg, S. & Dedek, K. (2019) Gap Junctions in A8 Amacrine Cells Are Made of Connexin36 but Are Differently Regulated Than Gap Junctions in AII Amacrine Cells. Front Mol Neurosci, 12, 99. PMC6489437

Yan, W., Laboulaye, M.A., Tran, N.M., Whitney, I.E., Benhar, I. & Sanes, J.R. (2020) Molecular identification of sixty-three amacrine cell types completes a mouse retinal cell atlas. bioRxiv, 2020.03.10.985770.

Zalutsky, R.A. & Miller, R.F. (1990) The physiology of substance P in the rabbit retina. J Neurosci, 10, 394-402. PMC6570149

Zeng, H. & Sanes, J.R. (2017) Neuronal cell-type classification: challenges, opportunities and the path forward. Nat Rev Neurosci, 18, 530–546.

Zhang, Y., Kim, I.J., Sanes, J.R. & Meister, M. (2012) The most numerous ganglion cell type of the mouse retina is a selective feature detector. Proc Natl Acad Sci U S A, 109, E2391-2398. PMC3437843

Zhu, Y., Xu, J., Hauswirth, W.W. & DeVries, S.H. (2014) Genetically targeted binary labeling of retinal neurons. J Neurosci, 34, 7845-7861. PMC4044247

